# Cellular reprogramming of human monocytes is regulated by time-dependent IL4 signalling and NCOR2

**DOI:** 10.1101/204180

**Authors:** Jil Sander, Susanne V. Schmidt, Branko Cirovic, Naomi McGovern, Olympia Papantonopoulou, Anna-Lena Hardt, Anna C. Aschenbrenner, Christoph Kreer, Kreer Quast, Alexander M. Xu, Lisa M. Schmidleithner, Heidi Theis, Thi Huong Lan Do, Hermi Rizal Bin Sumatoh, Mario A. R. Lauterbach, Jonas Schulte-Schrepping, Patrick Gunther, Jia Xue, Kevin Baßler, Thomas Ulas, Kathrin Klee, Stefanie Herresthal, Wolfgang Krebs, Bianca Martin, Eicke Latz, Kristian Händler, Michael Kraut, Waldemar Kolanus, Marc Beyer, Christine S. Falk, Bettina Wiegmann, Sven Burgdorf, Nicholas A. Melosh, Evan W. Newell, Florent Ginhoux, Andreas Schlitzer, Joachim L. Schultze

**Author notes:** Authors contributed equally (Jil Sander, Susanne V. Schmidt, Andreas Schlitzer, Joachim L. Schultze).

## Abstract

The clinical and therapeutic value of human *in vitro* generated monocyte-derived dendritic cell (moDC) and macrophages is well established. However, in line with recent findings regarding myeloid cell ontogeny and due to our limited understanding of their physiological counterparts, transcriptional regulation and heterogeneity, the full potential of these important cellular systems is still underestimated.

In this study, we use cutting edge high-dimensional analysis methods to better understand the transcriptional organization, phenotypic heterogeneity and functional differences between human *ex vivo* isolated and *in vitro* generated mononuclear phagocytes with the aim to better realize their full potential in the clinic.

We demonstrate that human monocytes activated by MCSF or GMCSF most closely resemble inflammatory macrophages identified *in vivo*, while IL4 signalling in the presence of GMCSF generates moDCs resembling inflammatory DCs *in vivo*, but not steady state cDC1 or cDC2. Moreover, these reprogramming regimes lead to activated monocytes that present with profoundly different transcriptomic, metabolic, phenotypic and functional profiles. Furthermore, we demonstrate that CD14^+^ monocytes are integrating multiple exogenous activation signals such as GMCSF and IL4 in a combinatorial and temporal fashion, resulting in a high-dimensional cellular continuum of reprogrammed monocytes dependent on the mode and timing of cytokine exposure. Utilizing nanostraw-based knockdown technology, we demonstrate that the IL4-dependent generation of moDCs relies on the induction, nuclear localization and function of the transcriptional regulator NCOR2.

Finally, we unravel unappreciated heterogeneity within the clinically moDCs population and propose a novel high-dimensional phenotyping strategy to better tailor clinical quality control strategies for patient need and culture conditions to enhance therapeutic outcome.

## Introduction

Recently, several transcriptomic, epigenetic, functional and fate-mapping studies established the identity of three different mononuclear cell lineages within the myeloid cell network, macrophages, dendritic cells (DC) and monocytes (Schlitzer et al., 2015a). During murine embryogenesis, different waves of progenitors colonize the developing tissues and establish the initial wave of tissue-resident macrophages, which are long lived and self-maintained in most tissues (Ginhoux and Guilliams, 2016; Ginhoux and Jung, 2014). Human and mouse DC can be separated into three distinct lineages, plasmacytoid DC, conventional DC (cDC) 1 and cDC2 (Merad et al., 2013; Satpathy et al., 2012; Schlitzer and Ginhoux, 2014). Both human and murine cDC1 and cDC2 have been shown to arise from specialized precursors within the bone marrow forming their functional specialization early during development, in contrast to the more plastic nature of macrophages (Breton et al., 2015; Ginhoux and Guilliams, 2016; Grajales-Reyes et al., 2015; Heidkamp et al., 2016; Lee et al., 2015; Perdiguero and Geissmann, 2016; Schlitzer et al., 2015b).

The third major component of the mouse and human mononuclear phagocyte network are monocytes. In the mouse, two phenotypically distinct subsets can be delineated, Ly6c^lo^ and Ly6c^high^ monocytes (Schlitzer et al., 2015a; Varol et al., 2015). In human peripheral blood, three different monocyte subsets can be identified by the expression of CD14 and CD16, in which CD14^+^CD16^−^ monocytes correspond to murine Ly6c^high^ monocytes and CD14^−^CD16+ monocytes are the counterpart of murine Ly6c^low^ monocytes (Auffray et al., 2009). Yet, little is known, whether this classification relates to functional specialization of distinct functional subsets.

Many studies suggest significant functional overlap between DCs, monocytes and tissue macrophages (Guilliams et al., 2014), including basic mechanisms such as phagocytosis (Fraser et al., 2009), anti-bacterial activity (Fehlings et al., 2012), antigen uptake, processing, and presentation (Cella et al., 1997; Dinter et al., 2014; Randolph et al., 2008), the capacity to activate adaptive immune cells (Toujas et al., 1997), cellular motility (Rossi et al., 2011), and activation programs (Frucht et al., 2000; Krutzik et al., 2005). The nature of this overlap has proven difficult to parse, but new knowledge concerning the distinct ontogeny of these cells has provided the opportunity to re-evaluate and elucidate the division of labor between DCs, monocytes and tissue macrophages.

Currently, assigning specific functions to each cellular entity within the mononuclear phagocyte system requires the use of *in vitro* systems to simplify complex cellular interactions and instead focus on basic molecular aspects. Murine bone marrow-derived DCs and macrophages cultures driven by granulocyte macrophage colony stimulating factor (GMCSF) or macrophage colony stimulating factor (MCSF) respectively, are frequently used to elucidate and assign molecular mechanisms of classical functions to subsets of mononuclear phagocytes. However these *in vitro* cultures have been shown to create a heterogeneous set of cells, making attribution of distinct cellular functions difficult (Helft et al., 2015). This conundrum highlights the need for a more detailed investigation of the cellular identity and the regulation therof in such *in vitro* cultures (Guilliams and van de Laar, 2015; Murray et al., 2014; Xue et al., 2014).

Sallusto *et al.* described the *in vitro* generation of human monocyte-derived DCs (moDCs) by culturing peripheral blood monocytes with GMCSF and IL4 (Sallusto and Lanzavecchia, 1994). Here, the term moDC was attributed to an activated monocyte population with DC-like functionality based on morphological and functional criteria. Similar phenotypes of functional convergence are observed in *in vitro* systems of human monocyte-derived macrophages driven by MCSF (moMΦ) (Akagawa et al., 2006) or GMCSF (Xue et al., 2014). Furthermore, systems biology-based definitions of mononuclear phagocyte function and nomenclature were established yielding insights about the identity, regulation and developmental origin of those cells (Guilliams and van de Laar, 2015; Murray et al., 2014; Xue et al., 2014). However, studies directly addressing their relationships to mononuclear phagocytes observed *in vivo* remain limited (Ohradanova-Repic et al., 2016).

Clarifying and assigning true functionality of *in vitro* generated monocyte-derived phagocytes to their *in vivo* counterparts and linking this knowledge to our most recent understanding of ontogeny is crucial considering the enormous interest in using *in vitro* generated mononuclear phagocyte derivatives for immunotherapeutic strategies in the clinic (https://clinicaltrials.gov/ct2/home). Therefore, the functional convergence, plasticity, and heterogeneity of monocyte-derived phagocytes highlighted above paired with the enormous clinical interest raises several important questions. Which part of the mononuclear cell compartment found *in vivo* do these different human *in vitro* culture systems represent? What is the difference between moMΦ further activated by IL4 (formerly described as ‘M2’ or better M(IL4) macrophages) and moDCs, which are similarly exposed to IL4, but with different time kinetics? Is there a temporal component in monocyte activation and how is this component integrated on the molecular level? And lastly, how heterogeneous are human *in vitro* monocyte cultures?

By applying transcriptome analysis combined with bioinformatics, the assessment of cellular phenotype and function, as well as nanostraw-delivered loss of function experiments, we elucidate the relationship of the widely used human moDCs and moMΦ systems to cells derived from the mononuclear phagocyte compartment *in vivo*. We also show that the cellular reprogramming of monocytes in these *in vitro* culture systems is multifaceted, integrating several time-dependent signals delivered by different growth factors and cytokines such as GMCSF and IL4. Additionally, we show that the time-dependent integration of IL4 during moDC differentiation is orchestrated by the nuclear receptor corepressor 2 (NCOR2). Finally, using mass cytometry, we show that clinically used moDCs are heterogeneous and consist of several different cellular entities.

## Results

### Cellular relationships of *in vitro* activated human monocyte-derived cells and *in vivo*mononuclear phagocytes

Human monocytes activated with MCSF have been used as models for human macrophages (Akagawa et al., 2006), while monocytes activated with GMCSF and IL4 were introduced as models for human DCs (Sallusto and Lanzavecchia, 1994). For clarity and in light of recent findings concerning DC, monocyte, and macrophage ontogeny (Ginhoux and Jung, 2014; Guilliams and van de Laar, 2015; Schlitzer et al., 2015a), we term activated monocytes according to their activation, e.g. monocytes activated with MCSF are named Mo-MCSF and monocytes activated for a specified duration (0-72h; 0-144h) with GMCSF and IL4 are Mo-GMCSF^IL4^. To establish the relationship and transcriptional similarity between *ex vivo* isolated cell subsets and activated monocytes, we compared blood derived CD14^+^ monocytes, CD1c^+^ DCs, CD141^+^ DCs (Haniffa et al., 2012), but also T- B- and NK-cells alongside CD45^+^lin^-^HLA-DR^high^ lung derived cells, to Mo-MCSF, Mo-GMCSF, Mo-GMCSF^IL4(0-72h)^, Mo-GMCSF^IL4(0-144h)^ using a genome-wide approach assessing whole transcriptomes (Figure S1A).

When reducing complexity of the data by principle component analysis (PCA), T-, B-, and NK-cells formed one of three larger clusters (green), all *ex vivo* isolated myeloid cells formed a second large cluster (red/yellow), and both these clusters were most distinct from a third cluster containing the four *ex vivo* polarized monocyte-derived cell populations (blue/purple,cyan,lilac, Figure 1A, B). We validated a three-cluster structure on the gene-level by hierarchical clustering (HC) of the 1,000 most variable genes (Figure 1C, **Table S1**). Pearson correlation coefficient matrix (PCCM) analysis further supported this cellular relationship model (Figure 1D), clearly showing that *in vitro* polarized monocytes are a transcriptomically separate entity compared to *ex vivo* isolated human peripheral blood lymphoid and myeloid cell types.

**Figure 1.**
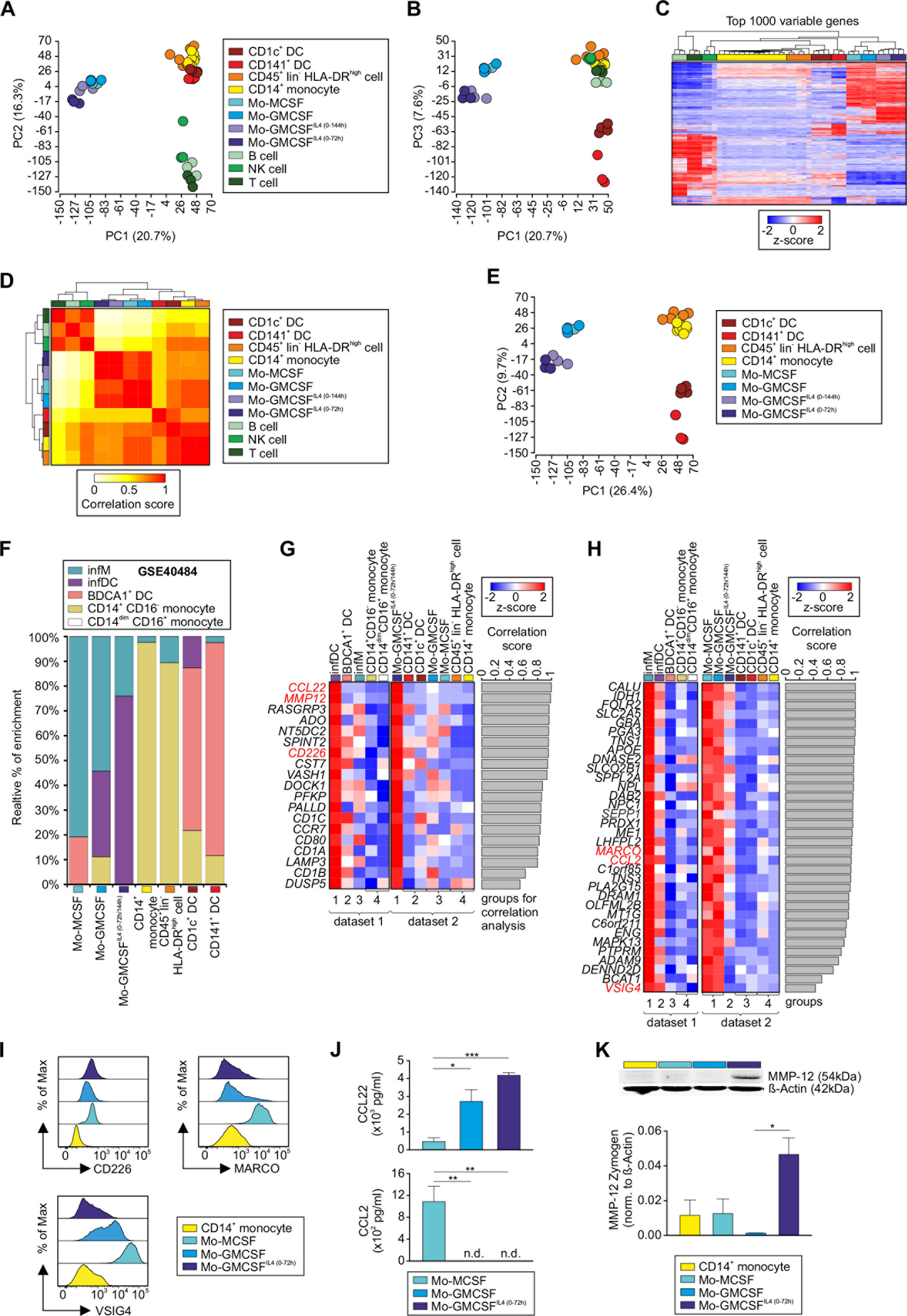
Relationship of *in vitro* activated monocyte-derived cells. **(A-B)** PCAs based on 21,250 present probes. Displayed are principal components (PCs) (**A**) 1 versus 2 and **(B)** 1 versus 3. **(C)** Heatmap of the top 1000 genes being most variable across the dataset. Log_2_-expression values were z-transformed and scaled (−2 (blue) to 2 (red)). **(D)** Heatmap visualizing Pearson correlation values (PCV) calculated pairwise between all cell types on the basis of the top 1000 most variable genes. **(E)** PCA based on 23,952 present probes. **(F)** Relative fractions of monocyte, BDCA1^+^ DC, infM and infDC gene signatures in CD14^+^ monocytes, different monocyte-derived cells and DCs. **(G-H)** Heatmaps of genes specifically expressed in **(G)** infDCs compared to infM, BDCA1^+^ DCs and monocytes (dataset 1), and in Mo-GMCSF^IL4(0-72h)^ cells compared to all other investigated monocyte-derived cells, CD14^+^ monocytes and DCs (dataset 2), or in **(H)** infM compared to infDCs, BDCA1^+^ DCs and monocytes (dataset 1), and in Mo-GMCSF and Mo-MCSF compared to Mo-GMCSF^IL4(0-72h)^, CD14^+^ monocytes and DCs (dataset 2). PCVs were calculated between the indicated group patterns of dataset 1 versus dataset 2 and are displayed as a barplot next to the heatmaps. Only genes with a correlation > 0.4 are displayed. Genes further analyzed in Figure 1 I-K are highlighted in red. Log_2_-expression values were z-transformed and scaled (−2 (blue) to 2 (red)). **(I)** Histograms show relative expression of CD226, MARCO and VSIG4 on CD14^+^ monocytes, Mo-MCSF, Mo-GMCSF and Mo-GMCSF^IL4(0-72h)^ analyzed by flow cytometry. Representative data from four different donors is shown. **(J)** Analysis of cell culture supernatants of Mo-MCSF, Mo-GMCSF and Mo-GMCSF^IL4(0-72h)^ for CCL22 and CCL2 using ELISA (n=3 with 2 technical replicates each, mean + SEM). Statistical significance was determined using one-way RM (repeated measures) ANOVA and Tukey’s method for multiple test correction, with \**p* < 0.05, \*\**p* < 0.01, and \*\*\**p* < 0.001. n.d. = not detected. **(K**) Relative quantification of MMP12 in CD14^+^ monocytes, Mo-MCSF, Mo-GMCSF and Mo-GMCSF^IL4(0-72h)^ using Western blot (n=3, mean + SEM). Statistical significance was determined using one-way ANOVA and Tukey's method for multiple test correction, with \**p* < 0.05.

To focus on the transcriptional analysis of mononuclear phagocyte subsets, we excluded T-, B- and NK cells from the previous dataset and used PCA to understand the transcriptional similarity between the remaining cell types (Figure 1E, **S1B, C)**. Interestingly, all *in vitro* generated cells cluster separately in comparison to blood myeloid cells found *in vivo*, with lung-derived CD45^+^lin^-^HLA-DR^high^ and CD14^+^ monocytes forming a separate cluster away from a distinct DC cluster as well as a separate cluster for Mo-MCSF, Mo-GMCSF, Mo-GMCSF^IL4(0-144h)^, Mo-GMCSF^IL4(0-72h)^. Additionally, all *in vitro* generated monocyte-derived subsets share a common set of differentially expressed genes in comparison to *ex vivo* isolated CD14^+^ monocytes (Figure S1D, E, **Table S1**), clearly supporting the transcriptional difference to homeostatic cells. Consequently, these findings raise the question which cells found *in vivo* are represented by the monocyte model systems.

Therefore, we assessed next if the *in vitro* polarization models of monocytes reflect the biology of human monocyte-derived inflammatory DCs (infDCs) and inflammatory macrophages (infM) *in vivo* (Segura et al., 2013). To this end, we utilized a previously published dataset (GSE40484, Affymetrix 1.1^ST^, (Segura et al., 2013)), generated a signature matrix representing infDCs, infM, BDCA1^+^ (CD1c) DCs, CD14^+^ CD16^−^ monocytes and CD14^dim^ CD16^+^ monocytes and assessed the relative enrichment of these signatures in our own dataset using linear support vector regression (Newman et al., 2015) (Figure 1F, **Table S1**). Mo-MCSF showed the highest enrichment of the infM-associated gene signature, while the infDC gene set was most highly enriched in Mo-GMCSF^IL4^, indeed suggesting that these *in vitro* polarization conditions reflect *in vivo* biology of infDCs and infM as previously shown for the mouse (Helft et al., 2015). Control gene sets derived from CD14^+^CD16^−^ monocytes were most highly enriched in *ex vivo* isolated CD14^+^ monocytes and lung-derived CD45^+^lin^−^HLA-DR^high^ cells, while the BDCA1^+^ DC signature was enriched in both *ex vivo* isolated myeloid DC subsets of our dataset. Gene Set Enrichment Analyses (GSEA) confirmed the transcriptional similarities between infM and Mo-MCSF but also Mo-GMCSF, as well as between infDC and Mo-GMCSF^IL4^ (Figure S1F). Based on those observations, we defined four (corresponding) groups in both datasets, describing comparable cell subsets (see Supplementary Methods). Then, we performed Pearson coefficient correlation analysis on the gene level by comparing the expression patterns in both datasets based on the four groups. We visualized the genes with the highest correlation scores between infDC and Mo-GMCSF^IL4 (0-72h/144h)^ (Figure 1G, **Table S1**) and between infM and both Mo-MCSF and Mo-GMCSF (Figure 1H, **Table S1**), which included several surface markers and secreted molecules (Figure S1G, H, I, J, **Table S1**). Clearly, many genes associated with activated DCs, such as *CCL22*, *MMP12*, *CD226*, and *CCR7*, were highly elevated in both infDC and Mo-GMCSF^IL4^ (Figure 1G), while typical macrophage genes including *MARCO*, *CCL2* and *VSIG4* were most highly expressed in infM, Mo-MCSF, and Mo-GMCSF (Figure 1H). Furthermore, differential regulation of CD226, MARCO, VSIG4, CCL2, CCL22 and MMP12 was also validated on the protein level, exemplifying the power of this computational approach (Figure 1I, J, K). Collectively, polarization of monocytes both *in vivo* (infDC, infM) (Segura et al., 2013) and *in vitro* leads to overlapping transcriptional reprogramming including many cell surface and effector molecules, which allows us to use these *in vitro* systems as a reductionist model to better understand the role of MCSF, GMCSF and IL4 during inflammation-induced monocyte activation.

### GMCSF + IL4 but not GMCSF or MCSF alone enforce a unique transcriptional signature in human CD14^+^ monocytes

Next, we were interested in better understanding similarities and differences in monocyte activation induced by the three model stimuli MCSF, GMCSF and IL4. Previous work suggested significant differences between Mo-MCSF and Mo-GMCSF (Lacey et al., 2012). However, these studies did not answer the overall relationship between all three activation conditions (Figure 2A). Using the well-established surface markers CD14, CD11b and CD209, we tried to delineate differences between Mo-MCSF, Mo-GMCSF and Mo-GMCSF^IL4(0-72h)^ (Figure S2A). This revealed that CD14 marked monocytes, Mo-MCSF and Mo-GMCSF, but not Mo-GMCSF^IL4(0-72h)^ as previously shown (Sallusto and Lanzavecchia, 1994). CD209 was exclusively expressed by Mo-GMCSF^IL4(0-72h)^. CD11b was indiscriminately expressed by all cell populations assessed. These results prompted us to elucidate the overall relationship between *ex vivo* isolated CD14^+^ monocytes, Mo-MCSF, Mo-GMCSF and Mo-GMCSF^IL4(0′72h)^ using a global transcriptomic approach (Figure S2B). PCA revealed a close relationship between Mo-MCSF and Mo-GMCSF, while monocytes and Mo-GMCSF^IL4(0-72h)^ were clearly distinct from the two former groups (Figure 2B). This relationship was further validated on gene level by hierarchical clustering (HC) of the 1,000 most variable genes revealing that Mo-MCSF and Mo-GMCSF formed mixed gene clusters (turquoise/light blue), marked by the high expression of macrophage related genes such as *CD81*, *VSIG4*, *SIGLEC1*, *MARCO* and *FPR3* (Figure 2C, **Table S2**). Contrary to this, monocytes and Mo-GMCSF^IL4(0-72h)^ cell populations formed separated gene clusters marked by the expression of key monocyte (*AHR*, *SELL*, *CLEC4D*) or DC-associated genes (*CD1C, ZBTB46*), respectively. PCCM analysis confirmed these findings (Figure S2C). Gene-level analysis only including the present surfaceome of *ex vivo* isolated CD14^+^ monocytes, Mo-MCSF, Mo-GMCSF and Mo-GMCSF^IL4(0-72h)^ revealed only a small number of differentially expressed transcripts for Mo-MCSF and Mo-GMCSF populations but a markedly different expression profile of surface markers related to Mo-GMCSF^IL4(0-72h)^ (Figure 2D). Additionally, profiling the expression of pattern recognition receptors (PRRs) expressed by *ex vivo* isolated CD14^+^ monocytes, Mo-MCSF, Mo-GMCSF and Mo-GMCSF^IL4(0-72h)^ revealed striking differences between these subsets (Figure S2D). Mo-MCSF, Mo-GMCSF and Mo-GMCSF^IL4(0-72h)^ markedly downregulated the mRNA levels of important components of the inflammasome signalling complex, such as *CASP1, NLRP1, 2* and *3*, but upregulated the intracellular PRR *NOD1. NOD2* expression was only maintained by Mo-GMCSF. Mo-GMCSF^IL4(0-72h)^ displayed a unique set of PRRs characterized by the high expression of *CD209* and *CLEC10A (CD301)* and the downregulation of Toll-like receptor (TLR) 7 and 5. To determine transcriptional differences, we performed co-expression network analysis based on the union of differentially expressed genes between monocytes and the three polarization conditions and mapped differential gene expression onto the network (Figure 2E, **Table S2**). Within the network topology, a large monocyte-related gene cluster was centrally placed surrounded by separate clusters for each of the three polarization conditions, further indicating that despite an overall very close relationship, Mo-MCSF and Mo-GMCSF are characterized by signal-specific subclusters of regulated genes (Figure 2E). Identification of differentially expressed genes between monocytes and Mo-MCSF and Mo-GMCSF further supported a close overall relationship, but also indicated significantly differently regulated genes in only one or the other condition (Figure S2E, F, **Table S2**). To investigate whether this would have functional implications, we performed a Gene Ontology Enrichment Analysis (GOEA) revealing that highly enriched terms in Mo-GMCSF are associated to immune response and regulation of protein metabolism, whereas most GO terms represented by Mo-MCSF are related to metabolism and G-protein coupled receptor signalling (Figure S2G, **Table S2**).

**Figure 2.**
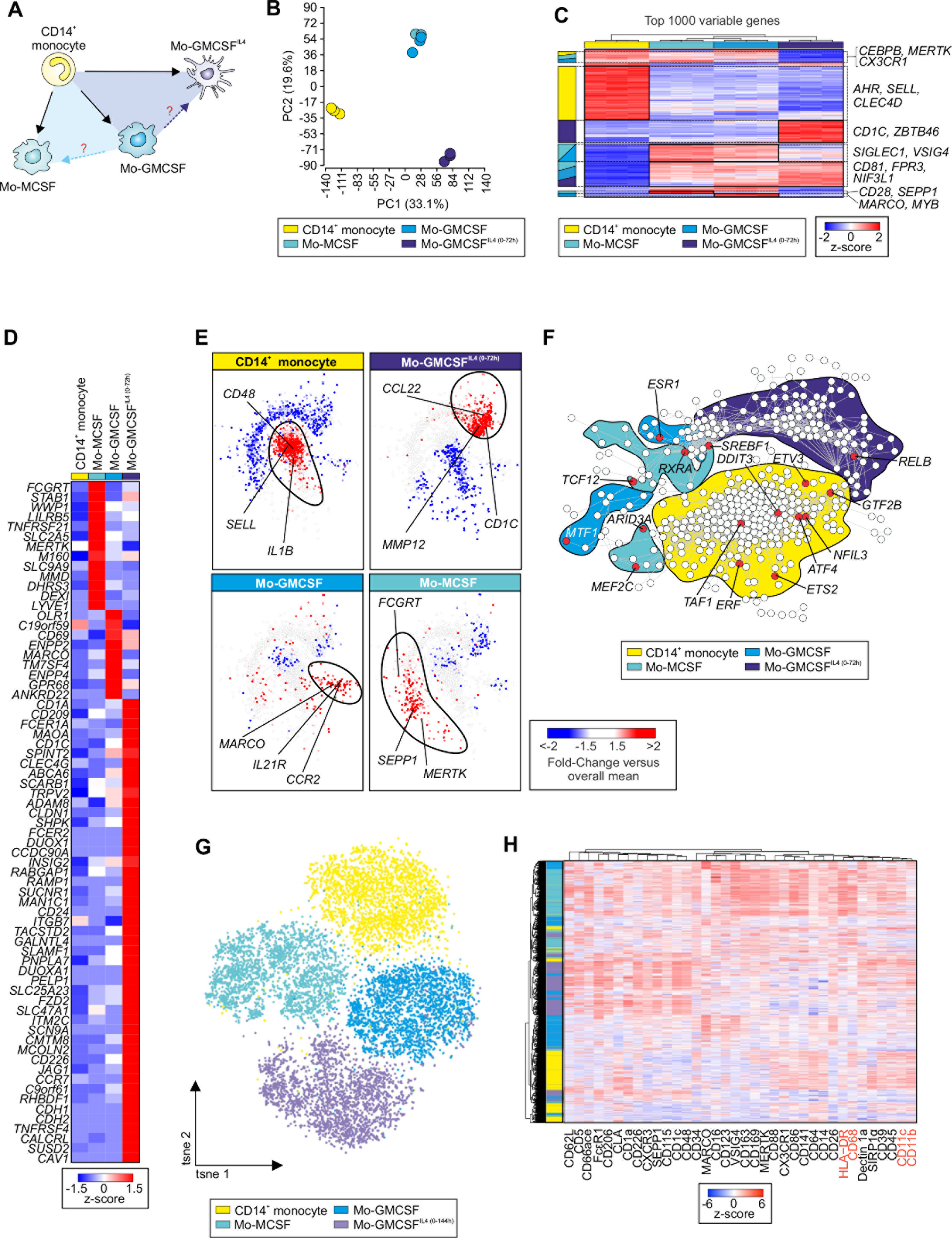
Mo-GMCSF^IL4^ are most distinct from Mo-MCSF and Mo-GMCSF. **(A)** Schema describing the biological questions addressed in Figure 2 and S2. (**B**) PCA based on 18,318 present probes. (**C**) Heatmap of the top 1000 genes being most variable across the dataset. Log_2_-expression values were z-transformed and scaled (−2 (blue) to 2 (red)). **(D**) Heatmap of genes being specifically expressed in a single out of the three monocyte-derived cell types, compared to CD14^+^ monocytes as well as to the other two monocyte-derived cell types. Log_2_-expression values were z-transformed and scaled (−1.5 (blue) to 1.5 (red)). (**E**) Co-expression networks based on the union of 2,086 differentially expressed genes (Fold-Change > 2 or < −2 and FDR-adjusted *p*-value < 0.05) between each of the three types of monocyte-derived cells compared to CD14^+^ monocytes. The Fold-Change of the respective cell type compared to the overall mean was mapped onto the networks and displayed in colors ranging from blue (negative Fold-Change) over white to red (positive Fold-Change). Based on the Fold-Change patterns, the networks were divided into four main clusters, each cluster representing one of the four cell types, respectively. (**F**) Co-expression network based on 411 TRs expressed in the dataset containing CD14^+^ monocytes, Mo-MCSF, Mo-GMCSF and Mo-GMCSF^IL4(0-72h)^. For each cell type, the Fold-Change of the respective cell type compared to the overall mean was mapped onto the network. Cell type-specific clusters of upregulated regulators were generated, indicated by the color-coded shadings behind the network. TRs highlighted in red were predicted to be unique master regulators for the corresponding cell type. The prediction was performed on all genes being highlighted in red (upregulated in the cell type compared to the overall mean with a Fold-Change > 1.5) in the corresponding cell type-specific cluster in **E**. Master regulators, which were identified for more than one cell type, were excluded. **(G)** *t*-SNE composite dimensions (tsne1 and 2) of CD14^+^ monocytes, Mo-MCSF, Mo-GMCSF and Mo-GMCSF^IL4(0-144h)^ analyzed by mass cytometry (3 intermixed donors shown). **(H)** Heatmap and hierarchical clustering of mean surface marker expression analyzed using mass cytometry on CD14^+^ monocytes, Mo-MCSF, Mo-GMCSF and Mo-GMCSF^IL4(0-144h)^. Normalized intensity values were z-transformed and scaled (−6 (blue) to 6 (red)). Color code depicts cluster assignment according to culture condition. Colour code as in **G**.

To identify the transcriptional regulators (TRs) involved in the generation of the monocyte-derived cell types, we predicted major upstream transcription factors (see Supplementary Methods) for each of the four condition-specific clusters identified in Figure 2E. We then generated a co-expression network of TRs expressed in the dataset and subsequently identified specific clusters of upregulated TRs for CD14^+^ monocytes (yellow), Mo-MCSF (turquoise), Mo-GMCSF (light blue) and Mo-GMCSF^IL4(0-72h)^ (dark blue) (Figure 2F, **Table S2**). Finally, we mapped the predicted master transcription factors onto the co-expression network and identified *NFIL3, ATF4* and *ETS2* among others to be specific regulators of CD14^+^ monocytes. *TCF12, MEF2C* and *ARID3A* specifically regulated the transcriptional identity of Mo-MCSF, whereas *ESR1, MTF1* and *SREBF1* were indicated to regulate the transcriptional makeup of Mo-GMCSF. Interestingly, *RELB*, which has been implicated to be important during mouse DC differentiation (Wu et al., 1998), was the only transcription factor predicted to be central to the formation of the transcriptional identity of Mo-GMCSF^IL4(0-72h)^, further highlighting the uniqueness of the transcriptional reprogramming induced by GMCSF and IL4.

Since the usage of traditional surface markers, such as CD14, CD11b and CD209 to discriminate *in vitro* polarized monocyte subsets was not very informative, we designed a comprehensive mass cytometry panel incorporating established and novel protein markers defined by our global transcriptomic approach and compared *ex vivo* isolated blood CD14^+^ monocytes to *in vitro* MCSF, GMCSF, GMCSF + IL4 polarized CD14^+^ monocytes. Dimensionality reduction using *t*-distributed neighbour embedding (*t*-SNE) (Maaten and Hinton, 2008) of the CD45^+^lin^−^HLA-DR^+^cell fraction comparing CD14^+^ monocytes, Mo-MCSF, Mo-GMCSF and Mo-GMCSF^IL4(0-144h)^ revealed donor-independent clustering into four different cellular subsets (Figure 2G, H, **S2H**). Overlaying their polarization history on the *t*-SNE topology revealed that the four identified clusters corresponded to the four different polarization conditions, validating the differences found within the global transcriptional data (Figure 2B). Interestingly, commonly used markers for the delineation of monocytes, macrophages and DCs, such as CD11b, CD68, CD11c and HLA-DR were expressed uniformly across all four clusters, showing that only a high-dimensional phenotyping approach enables robust detection of polarized subsets across all four polarization conditions (Figure 2H). CD14^+^ monocytes were characterized by a high expression of CLA and CD64, whereas Mo-GMCSF displayed a high expression of MARCO. VSIG4 was commonly expressed by Mo-GMCSF and Mo-MCSF, whereas Mo-MCSF specifically expressed high levels of the macrophage related proteins CD163, CD169 and MERTK. The Mo-GMCSF^IL4(0-144h)^ cluster was characterized by specific expression of SEPP1, FcsR1, CD1c and CD48. Taken together, mass cytometry enabled us to identify novel, transcriptionally validated markers, facilitating separation of different transcriptomic entities on the protein level.

### Functional properties of *in vitro* polarized monocytes

To understand how the observed transcriptomic and phenotypic differences of Mo-MCSF, Mo-GMCSF and Mo-GMCSF^IL4^ translate to the functional level, we assessed their ability to phagocytize, to secrete cytokines in response to PRR stimulation, their motility and their metabolic profile.

First, we measured receptor-mediated uptake of GFP-labelled yeast or YG beads. Mo-MCSF, Mo-GMCSF and Mo-GMCSF^IL4(0-72h)^ were equally able to phagocytose GFP^+^ yeast buds after 1 hour of incubation indicating that there is no differential induction of receptors and signalling pathways essentially involved in yeast uptake (Figure 3A, **S3A**). Mo-MCSF displayed an up to 12 times enhanced uptake of YG beads in comparison to Mo-GMCSF^IL4(0-72h)^ (Figure 3B, **S3B**). Therefore, despite their close relationship on the transcriptional level, these data suggest that MCSF activation but not GMCSF drives an overall phagocytic capacity in monocytes. When assessing cell motility (Figure 3C, **S3C, D**), Mo-MCSF, Mo-GMCSF and Mo-GMCSF^IL4(0-72h)^ cells showed very little, intermediate and high level motility, respectively. Accumulated distance and velocity analysis corroborated these findings suggesting that migratory capacity is linked to GMCSF activation, but not MCSF and is potentiated by IL4 signalling (Figure 3C, **S3C, D**). Metabolically, Mo-MCSF and Mo-GMCSF show a similar rate of oxidative phosphorylation (OXPHOS), extracellular acidification (ECAR), ATP production and glycolytic capacity (Figure 3D-E, **S3E-I**). Mo-GMCSF^IL4(0-72h)^ however display a statistically significant increase of OXPHOS, ATP production and glycolytic capacity alongside an elevated maximal respiration capacity indicative of increased energetic fitness alongside higher metabolic flexibility induced by IL4-specific reprogramming (Figure 3D-E, **S3E-I**).

**Figure 3.**
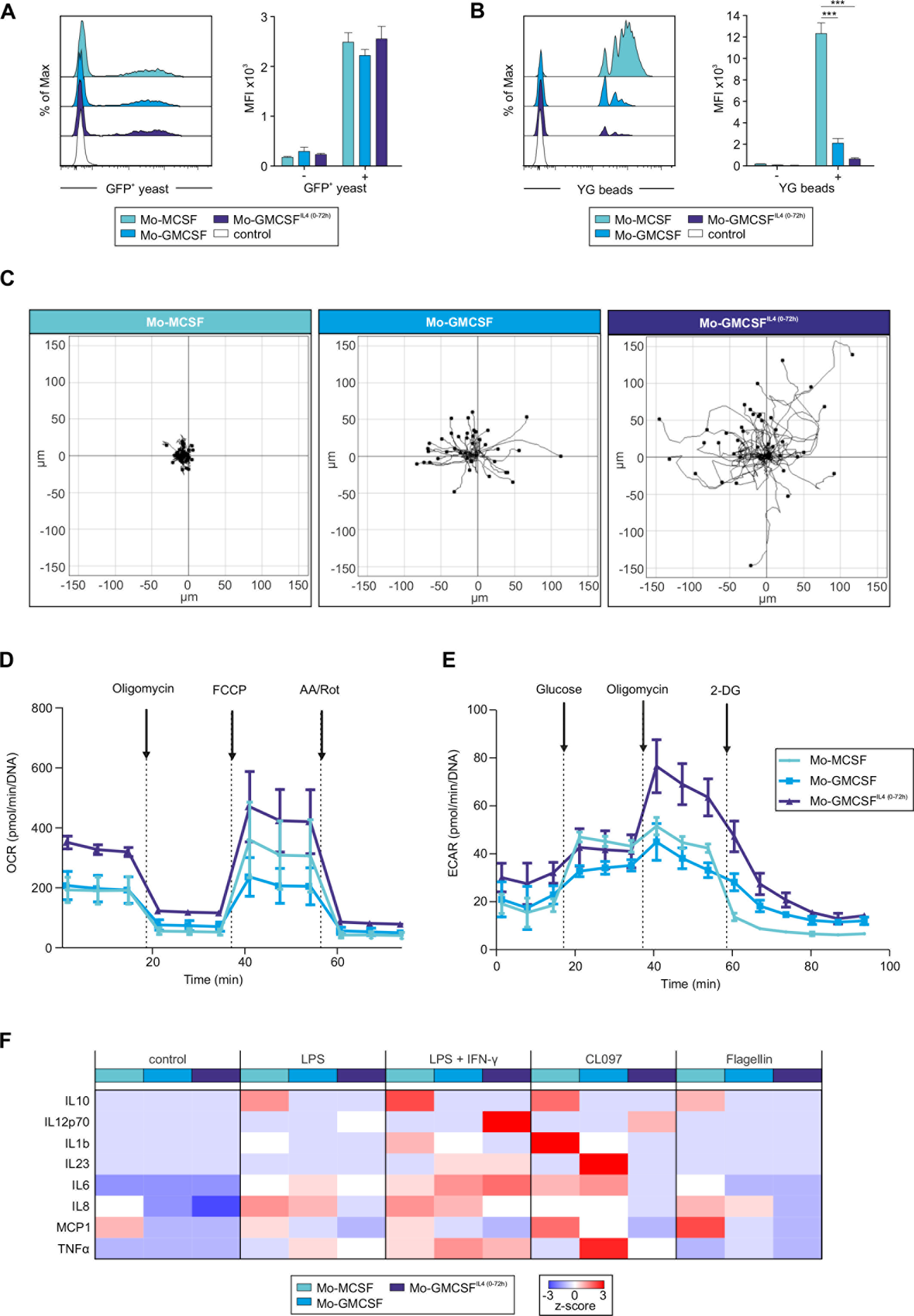
Prediction of activated monocyte functionality. **(A)** Flow cytometric analysis of Mo-MCSF, Mo-GMCSF and Mo-GMCSF^IL4(0-72h)^ after 1h of incubation with GFP-expressing yeast (histogram: 1 representative of 3 replicates, bar plot n=3, mean + SEM). Statistical significance between cell types after 1h of incubation was determined using one-way RM ANOVA, with (*p* > 0.05). **(B)** Flow cytometric analysis of Mo-MCSF, Mo-GMCSF and Mo-GMCSF^IL4(0-72h)^ after 4hrs of incubation with YG beads (n=5-6, mean + SEM). Statistical significance between cell types after 4h of incubation was determined using one-way RM ANOVA and Tukey’s method for multiple test correction, with \*\*\**p* < 0.001. **(C)** Migration tracks of Mo-MCSF, Mo-GMCSF and Mo-GMCSF^IL4(0-72h)^ migrating on a surface coated with fibronectin for 3hrs. Results show one representative experiment out of three. **(D)** OCR of Mo-MCSF, Mo-GMCSF and Mo-GMCSF^IL4(0-72h)^, followed by sequential addition of oligomycin, FCCP, and Rot/AA. **(E)** ECAR of Mo-MCSF, Mo-GMCSF and Mo-GMCSF^IL4(0-72h)^, followed by sequential addition of glucose, oligomycin and 2-Deoxyglucose. **(F)** Heatmap displaying mean secreted cytokine concentrations of up to four donors. Expression values were z-transformed and scaled (−3 (blue) to 3 (red)). Raw data can be found in **Table S3**.

Linking these functional data concerning phagocytosis, migration and metabolism back to the global transcriptomic approach, we identified key genes involved in the regulation of these processes. *RAB10* (Cardoso et al., 2010), *MSR1* (Bonilla et al., 2013) and *DAB2* (Tumbarello et al., 2013) among others have been implicated to play a pivotal role in the regulation of phagocytosis in immune cells and subsequent antigen processing. Interestingly, these genes alongside other regulators of phagocytosis, such as *RAPH1, RILPL2, TNS1* and *SCARB2*, are markedly upregulated on the gene level in Mo-MCSF (Figure S3J, **Table S3**). Similarly, regulators of migration and cell motility such as Lymphotoxin β (*LTB*) (Yu et al., 2002), *CCL13* (Stellato et al., 1997), *CCL22* (Godiska et al., 1997) and *ASAP1* (Curtis et al., 2015) were upregulated in Mo-GMCSF^IL4(0-72h)^ connecting transcriptomic identity of this cell population clearly with its unique functional properties (Figure S3K, **Table S3**). Additionally, genes involved in the regulation of glycolysis such as *PFKL* and *PFKP* were highly upregulated in Mo-GMCSF^IL4(0-72h)^ corresponding to the higher glycolytic capacity of these cells (Figure S3L, **Table S3**). Furthermore *UQCRC1, SDHA, ATP5D, COX10, ATP5I* genes of the respiratory chain and *IDH3G*, a molecule involved in the TCA cycle were highly upregulated in Mo-GMCSF^IL4(0-72h)^, further corroborating that the functional changes also transcend to the transcriptional level.

Finally, we stimulated Mo-MCSF, Mo-GMCSF and Mo-GMCSF^IL4(0-72h)^ with LPS, LPS + interferon γ (IFN-γ), CL097 or Flagellin to evaluate their potential to secrete cytokines upon PRR ligation (Figure 3F, **Table S3**). Interestingly, the major immunoregulatory cytokines IL10, IL1B and MCP1were secreted only by Mo-MCSF upon activation with either of the four stimuli. This further demonstrates their similarities to *in vivo* infM, which have very plastic cytokine secretion properties. Conversely, IL12p70 was only secreted by Mo-GMCSF^IL4(0-72h)^ upon stimulation with either LPS, LPS + IFN-γ and CL097, indicating that these cells indeed possess functional overlap with DCs regarding the induction of Th1 T-cell responses through the secretion of IL12p70. IL23, a major driver of inflammatory reactions being essential for the induction of Th17 responses was only produced by cells which have been reprogrammed by GMCSF. Both Mo-GMCSF and Mo-GMCSF^IL4(0-72h)^ were able to produce IL23 following LPS + IFN-γ and CL097 stimulation, respectively. This is in line with their similarity to ovarian cancer induced infDCs, which have been shown to have the superior capacity to induce Th17 immunity (Segura et al., 2013).

### IL4 regulates transcriptomic and functional polarization of moDCs and monocyte-derived “M2-like” macrophages

Presuming that IL4 is inducing a functional convergent phenotype between monocytes and DCs, we next asked whether Mo-GMCSF^IL4(0-144h)^ are distinct from what was previously described as M2 macrophages, better described as monocyte-derived macrophages further activated by IL4, termed here Mo-GMCSF^IL4(72-144h)^. To reduce variables to a minimum, we generated Mo-GMCSF^IL4(72-144h)^ with GMCSF, so that the only differences to Mo-GMCSF^IL4(0-144h)^ cells were the onset and the overall time of IL4 exposure. As controls, we also included monocytes polarized for only 3 days with either GMCSF (Mo-GMCSF^IL4(0h)^) or GMCSF+IL4 (Mo-GMCSF^IL4(0-72h)^) (Figure 4A). Globally, as determined by PCA, Mo-GMCSF^IL4(72-144h)^ were surprisingly distinct from (Mo-GMCSF^IL4(0-72), (0-144h)^) irrespective whether Mo-GMCSF^IL4^ were sampled after 72h or 144h of activation (Figure 4B, **S4A, B**). These findings were further corroborated by co-expression network analysis (Figure 4C), HC using the most variable genes (Figure 4D, **Table S4**), and PCCM analysis (Figure S4C). When assessing immune-phenotypes, a surprising finding was the similarly high expression of CD23 (Figure S4D), a marker formerly associated with Mo-GMCSF^IL4(72-144h)^ (Mantovani et al., 2002), and comparably high expression of CD209 (Figure S4E), which has been linked to monocytes with DC functionality (Geijtenbeek et al., 2000). There was no significant difference in the expression of MMP-12 (Figure S4F, G), but in the release of CCL22 (Figure S4H). In contrast, Mo-GMCSF^IL4(0-144h)^ showed higher uptake of yeast within 60 minutes of exposure time (Figure 4E), but similar levels of phagocytosis of YG beads (Figure S4I) when compared to Mo-GMCSF^IL4(72-144h)^. Mo-GMCSF^IL4 (0-144h)^ were also more motile than Mo-GMCSF^IL4(72-144h)^ (Figure 4F), with both statistically higher accumulated distance (Figure S4J) and velocity (Figure S4K). Analysis of the metabolic parameters of Mo-GMCSF^IL4(0-144h)^ and Mo-GMCSF^IL4(72-144h)^ revealed no differences in their rate of OXPHOS, ATP production and glycolysis (data not shown). Collectively, these data strongly suggest that integration of the IL4 signalling is regulated in a time-dependent manner and represents a critical checkpoint for monocyte reprogramming.

**Figure 4.**
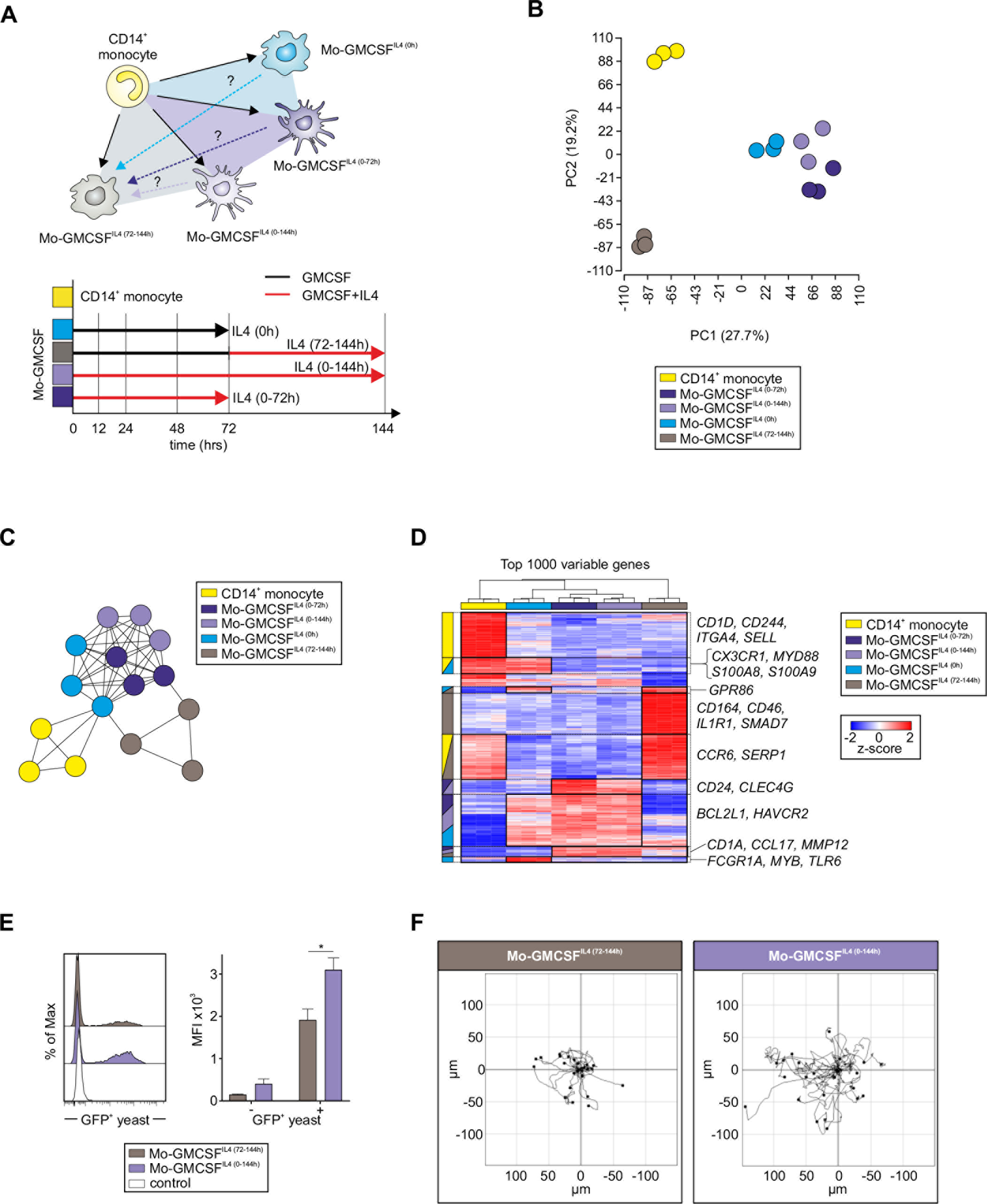
Mo-GMCSF^IL4(0-144h)^ differ from Mo-GMCSF^IL4(72-144h)^ monocyte-derived cells. **(A)** Schema describing the biological questions addressed in Figure 4 and S4. **(B)** PCA based on 18,857 present probes. **(C)** Co-expression network describing the relationships between all samples of the dataset containing CD14^+^ monocytes and four types of monocyte-derived cells based on 13,691 present genes. **(D)** Heatmap of the top 1000 genes being most variable across the dataset. Log_2_-expression values were z-transformed and scaled (−2 (blue) to 2 (red)). Genes were grouped together (black boxes), dependent on in which cell type(s) they appeared as highly expressed. The corresponding group-related cell types are highlighted on the left side next to the heatmap (color coded). Important genes of each cluster are depicted on the right side next to the heatmap. **(E)** Flow cytometric analysis of Mo-GMCSF^IL4(72-144h)^ and Mo-GMCSF^IL4(0-144h)^ after 1h of incubation with GFP-expressing yeast (n=4-6, mean + SEM). Statistical significance between cell types after 1h of incubation was determined using Student’s t-test with \**p* < 0.05. **(F)** Migration tracks of Mo-GMCSF^IL4(72-144h)^ and Mo-GMCSF^IL4(0-144h)^ migrating on a surface coated with fibronectin for 3hrs. Results show one representative experiment out of three.

### Timing of IL4 stimulation determines transcriptional regulation in activated monocytes

The experiments presented in Figure 4 allowed two alternative explanations: a dichotomous model with monocytes differentiating into monocytes with DC or macrophage functionality, or alternatively, a continuum model that integrates time of exposure suggesting significant plasticity of monocyte-derived cells. To determine which of these models described the influence of IL4 on monocytes best, we performed a time kinetics experiment, in which IL4 was added at the start of the culture (Mo-GMCSF^IL4(0-144h)^), or 12 (Mo-GMCSF^IL4(12-144h)^), 24 (Mo-GMCSF^IL4(24-144h)^), 48 (Mo-GMCSF^IL4(48-144h)^), or 72 hours (Mo-GMCSF^IL4(72-144h)^) after initiation of activation with GMCSF (Figure 5A). Whole transcriptomes were assessed after 144 hours and Mo-GMCSF^IL4(0h)^ and Mo-GMCSF^IL4(0-72h)^ were used as further controls. When assessing the expression of CD14 (Figure 5B, **S5A**) and CD209 (Figure 5C, **S5B**), we observed a dichotomous distribution for cells activated with IL4 being CD14^low^ CD209^high^ while cells activated only by GMCSF were CD14^+^CD209^−/low^. In contrast, global analysis on the transcriptional level revealed a different model (Figure S5C). Applying PCA, a gradual ordering of samples formed corresponding to the exposure time to IL4, indicating a gradual commitment to a polarization state along a continuum of plasticity (Figure 5D). HC of the 1000 most variable genes showed a similar order of the samples (Figure 5E, **Table S5**). More importantly, this analysis suggested a gradual change in gene expression dependent on the time of IL4 exposure which was further corroborated by SOM clustering (Figure 5E). Next, we applied co-expression network analysis and mapped gene expression information for each time point onto the network (Figure 5F, **Table S5**). This densely populated network was characterized by two major clusters, with one being characterized by genes elevated in monocytes not exposed to IL4 (0h, red: up-regulated; blue: down-regulated). In contrast, for each time point of IL4 exposure, a different set of genes within the other major cluster was most significantly elevated (Figure 5G). Adjacent time points even showed partially overlapping gene sets strongly suggesting a plastic continuum integrating IL4 signalling over time, arguing against the dichotomous model of polarization.

**Figure 5.**
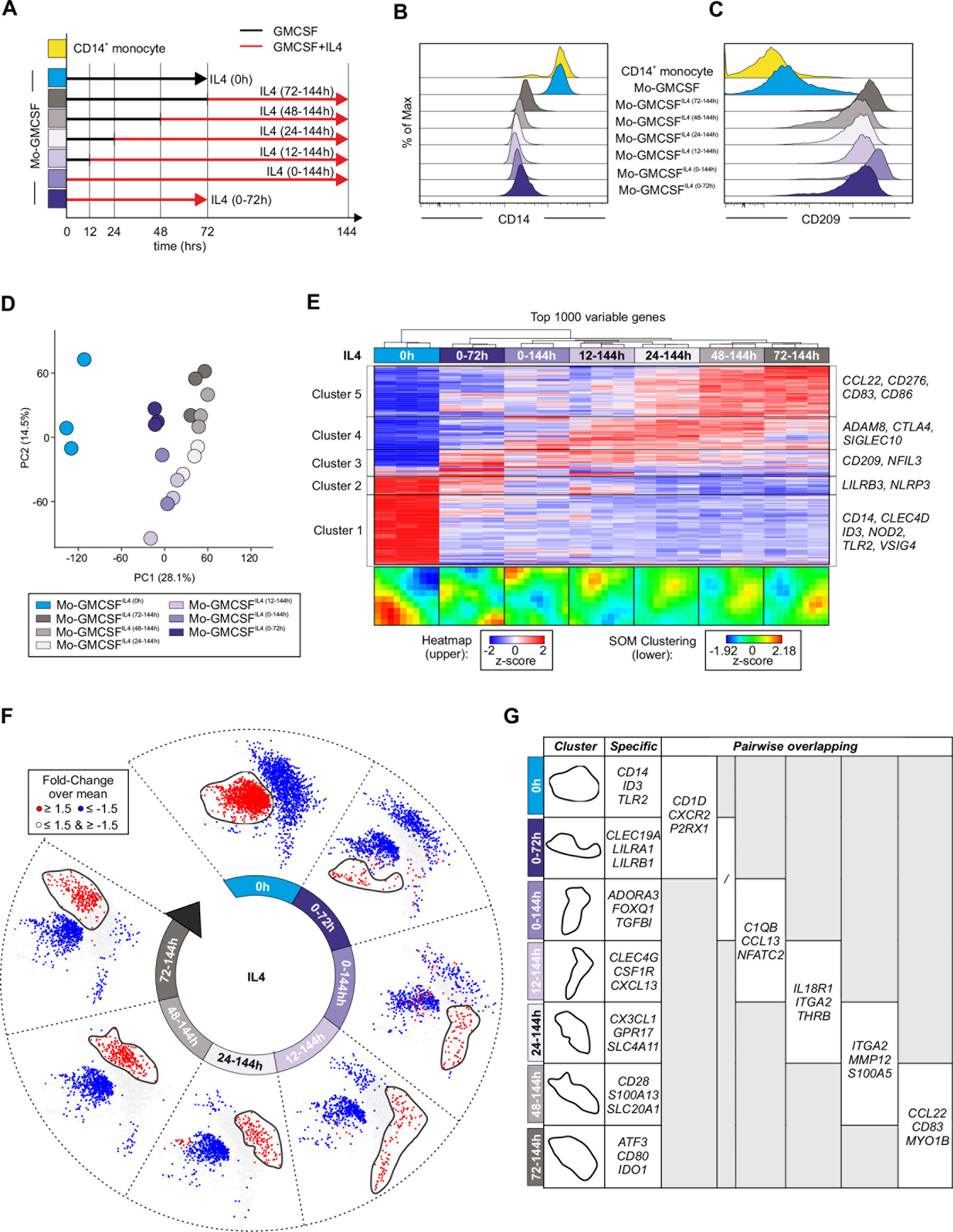
Timing of IL4 determines transcriptional regulation in activated monocytes. **(A)** Schema describing the approach of the time kinetic experiment. **(B-C)** Histograms show relative expression of CD14 and CD209 on the depicted culture timings analyzed by flow cytometry. **(D)** PCA based on 12,794 present genes. **(E)** Heatmap of the top 1000 genes being most variable across the dataset. Log2-expression values were z-transformed and scaled (−2 (blue) to 2 (red)). Below SOM-clustering, determined based on the expression profiles of the 12,794 present genes across all cell types. **(F)** Co-expression networks based on the union of 2,775 genes being differentially expressed (Fold-Change > 1.5 or < −1.5 and FDR-corrected p-value < 0.05) between each monocyte-derived cell type activated by IL4 compared to Mo-GMCSF. For each cell type, the Fold-Change of the respective cell type compared to the overall mean was mapped onto the networks and displayed in blue (Fold-Change <= 1.5) or red (Fold-Change >= 1.5). **(G)** Examples of genes located in the condition-related clusters depicted in **F** and in the first column, having a Fold-Change >= 1.5 for the corresponding condition. Genes listed in the second column are condition-specific, genes displayed in the following columns are shared between the clusters of two consecutive time points.

Collectively, these data suggest that monocytes are reprogrammed by IL4 signalling over time along a continuum with Mo-GMCSF^IL4(0h)^ and Mo-GMCSF^IL4(72-144h)^ being at the extreme ends.

### NCOR2 is a transcriptional regulator of Mo-GMCSF^IL4^

To better understand how IL4 enforces this unique transcriptional program in Mo-GMCSF^IL4(0-72h)^, we generated a co-expression network by first extracting TRs from the transcriptome data expressed in either monocytes, Mo-GMCSF or monocytes activated with GMCSF and different durations of IL4 and used them as bait for building a co-expression network of TRs (Figure 6A, B). Secondly, TRs with differential expression in Mo-GMCSF^IL4(0-72h)^ in comparison to the other cell types were filtered and thirdly, the remaining TRs were ranked based on absolute expression levels (Figure 6B, C, red: upregulated, blue: downregulated, **Table S6**). We identified seven TRs fulfilling these criteria with NCOR2 showing the highest expression of these TRs (Figure 6C). Analysis of NCOR2 protein expression revealed an enrichment of NCOR2 in the nucleus of Mo-GMCSF^IL4(0-72h)^ but not in Mo-GMCSF or monocytes (Figure 6D). Using nanostraw technology (Figure S6B), which allows viral- or liposome free siRNA delivery (Xu et al., 2014), we introduced anti-NCOR2 siRNAs for the last 24 hours of the Mo-GMCSF^IL4(0-72h)^ activation program. After siRNA incubation, mean NCOR2 mRNA levels in Mo-GMCSF^IL4(0-72h)^ were reduced to 65% relative to the control (Figure S6C), which reflects effective downregulation of NCOR2 transcription considering its long half-life of more than 24 hours (Raghavan et al., 2002). Interestingly, siRNA knockdown of NCOR2 in Mo-GMCSF^IL4(0-72h)^ also reduced the levels of CD209 mRNA (Figure S6D) and protein (Figure S6E). To understand the global impact of NCOR2 on the transcriptional program of Mo-GMCSF^IL4(0-72h)^ and the regulation of Mo-GMCSF^IL4 (0-72h)^ transcriptional identity by NCOR2, we performed a global transcriptome analysis of anti-NCOR2 siRNA-treated Mo-GMCSF^IL4(0-72h)^ versus scrambled siRNA-treated Mo-GMCSF^IL4(0-72h)^ (Figure S6F, G). We identified 1,834 variable genes after knockdown of NCOR2 (Figure S6H, **Table S6**). To further classify NCOR2 genes, we defined an IL4 signature based on three previously described datasets (GSE13762, GSE35304, GSE32164) with a total of 457 signature genes induced and 498 genes repressed by IL4 (Figures S6F). Overlaying these signature genes onto an expression plot comparing the previously identified variable genes between control and NCOR2 knockdown samples demonstrates that a large majority of the genes regulated by NCOR2 are IL4 signature genes (Figure 6E, **Table S6**). Taken together, this data establishes NCOR2 as a key regulator involved in the reprogramming of monocytes by IL4 enforcing their unique transcriptional and functional profile.

**Figure 6.**
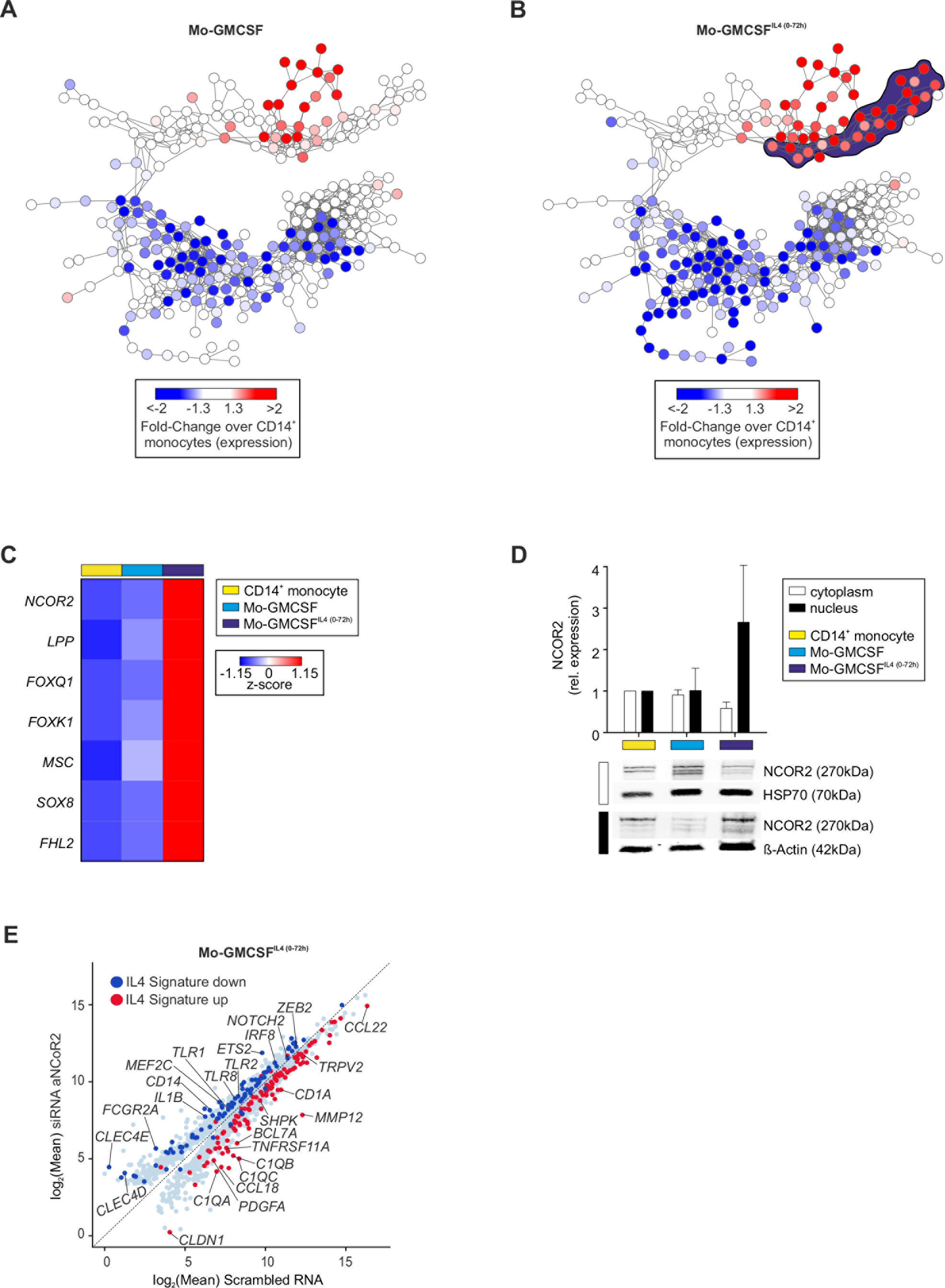
NCOR2 is a transcriptional regulator of Mo-GMCSF^IL4(0-72/144h)^. **(A-B)** Co-expression networks based on 267 TRs being expressed in CD14^+^ monocytes and four types of monocyte-derived cells. For **(A)** Mo-GMCSF and **(B)** Mo-GMCSF^IL4(0-72h)^, the fold-change compared to CD14^+^ monocytes was mapped onto the network. According to the resulting patterns, a Mo-GMCSF^IL4(0-72h)^-specific cluster of upregulated regulators was generated, which is indicated by the color-coded shading (dark blue) behind the network. Upregulated regulators in Mo-GMCSF^IL4(0-72h)^ (as determined in **C**) within the network are labeled in black. **(C)** Heatmap of TRs, which were identified to be specifically upregulated in Mo-GMCSF^IL4(0-72h)^ and Mo-GMCSF^IL4(0-144h)^ cells compared to CD14^+^ monocytes and Mo-GMCSF. Log_2_-expression values were z-transformed and scaled (−1.15 (blue) to 1.15 (red)). **(D)** Relative quantification of NCOR2 protein in cytoplasm and nucleus of CD14^+^ monocytes, Mo-GMCSF and Mo-GMCSF^IL4(0-72h)^ using Western blot alongside HSP70 and β-actin (n=3, mean + SEM). **(E)** Scatterplot of 1,834 variable genes across the dataset containing cells with siRNA αNCoR2 (y-axis) and control cells with scrambled RNA (x-axis). Displayed are log_2_-mean expression values. Highlighted genes were determined (as described in Figure S6F) to be either induced (red) or repressed (blue) by IL4 in external datasets.

### Mass cytometry analysis identifies unappreciated phenotypic heterogeneity in clinically relevant Mo-GMCSF^IL4(0-144h)^ cultures

Albeit high-dimensional protein analysis by mass cytometry clearly distinguished the four cell populations, namely monocytes, Mo-MCSF, Mo-GMCSF, and Mo-GMCSF^IL4(0-144h)^ (Figure 2G), we recognized a subcluster structure within the different cell subsets, which might indicate further unappreciated phenotypic heterogeneity within *in vitro* cultured cells. After complexity reduction, we used the Rphenograph package (Chen et al., 2016; Levine et al., 2015) to cluster the dataset and devise novel subclasses of cells within the identified clusters (Figure 7A, B). As expected, the top cluster corresponding to CD14^+^ monocytes did not reveal any additional heterogeneity for the tested markers and presented with a homogenous expression of the known monocyte markers CD14, CD11b, CX3CR1, alongside low expression of newly identified markers for monocyte reprogramming such as VSIG4, MARCO and SEPP1. Mo-MCSF revealed four subpopulations (cluster 2, 5, 10, 11), which were characterized by the co-expression of the tissue macrophage related markers MERTK, CD64, CD169 and CD163 and showed variable expression of L-selectin (CD62L, low in cluster 2, 11) and CD26 (low in cluster 2), indicating a different migration and maturation status (Figure 7B). High-dimensional analysis of Mo-GMCSF revealed two phenotypically different subclusters within this polarized monocyte populations (cluster 7 & 8). Both clusters uniformly expressed CD14, CD68 and CD206. Albeit CD64 was expressed by clusters 7 and 8, other macrophage-related proteins, such as MERTK, CD169 and CD163, were not, further corroborating their difference from Mo-MCSF on the phenotypic level. Cluster 7additionally expressed the activation associated molecules FcsR1 and CD1a, supporting a more activated status of this subpopulation of Mo-GMCSF. M-GMCSF^IL4(0-144h)^ showed a similar degree of heterogeneity as Mo-MCSF, however with more pronounced phenotypic differences (cluster 1, 3, 4, 9). To understand these differences in more detail, we isolated Mo-GMCSF^IL4(0-144h)^ cells out of the dataset and analyzed these cells alone using *t*-SNE dimensionality reduction and the Rphenograph clustering algorithm (Figure 7C-E). Rphenograph revealed 11 phenotypically different subpopulations within the population polarized by GMCSF and IL4 (Figure 7C, D). Common markers of moDC such as CD1c, CD226, CD48 and CD11c were uniformly expressed across all 11 subpopulations. Interestingly, the biggest differences across all different subpopulations were seen in the expression of activation and antigen presentation-associated molecules, such as CD1a, HLA-DR, FcsR1, CD62L and CD86. Expression of these markers differed across the subpopulations with clusters 6, 1, 10, 11 showing high expression of CD1a and CD62L, whereas only cluster 5 and 2 expressed high amounts of the important co-stimulatory molecule CD86. Of note, also the expression of CD11b, a marker widely used to analyze the purity of these cells in the clinic, varied significantly across the examined GMCSF + IL4 population, exemplifying the urgent need for a higher dimensional phenotyping to improve purity and therapeutic outcome when using these reprogrammed monocytes therapeutically (Figure 7E).

**Figure 7.**
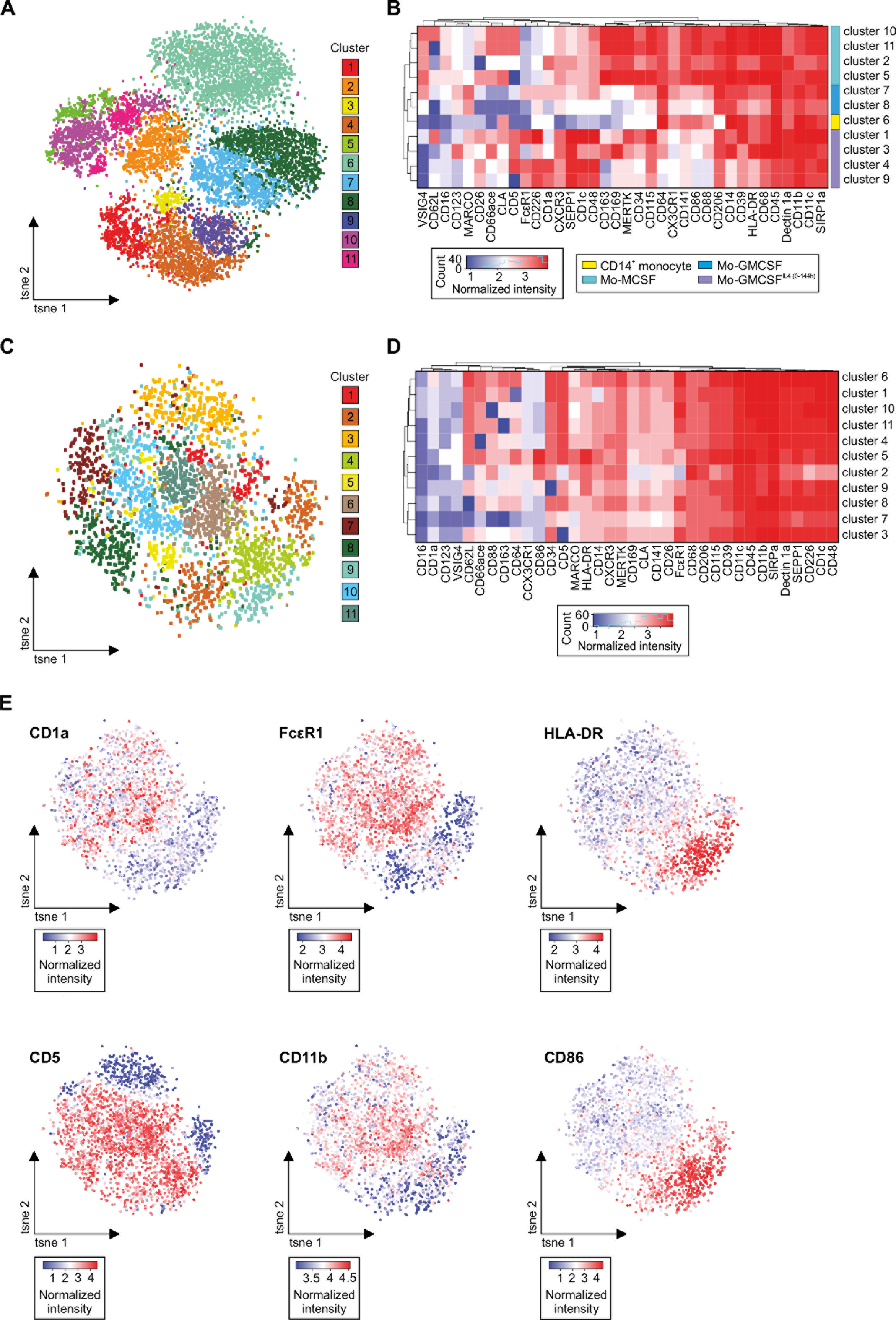
Mass cytometry analysis identifies unappreciated phenotypic heterogeneity in clinically relevant Mo-GMCSF^IL4(0-144h)^ cultures. **(A)** Phenograph analysis of CD14^+^ monocytes, Mo-MCSF, Mo-GMCSF and Mo-GMCSF^IL4(0-144h)^ based on mass cytometry expression data derived from three donors (as presented in Figure 2G) including 36 myeloid-related surface markers. Affiliation of individual cells to the identified clusters is indicated by color coding and visualized in a *t*-SNE plot. **(B)** Heatmap and hierarchical clustering of mean surface marker expression of the 11 individual clusters identified in (**A**). Reprogramming conditions matching the clusters according to A and Figure 2G are indicated on the right. **(C)** Mass cytometry analysis focusing on 3500 Mo-GMCSF^IL4(0-144h)^ cells derived from one representative donor analyzed using Phenograph and visualized in a *t*-SNE plot. **(D)** Heatmap and hierarchical clustering of mean surface marker expression of the 11 individual clusters identified in (C). **(E)** Feature plot display of the expression of selected surface markers (CD1a, FcεR1, HLA-DR, CD5, CD11b, CD86) in Mo-GMCSF^IL4(0-144h)^ cells overlaid onto the *t*-SNE plot.

## Discussion

Human CD14^+^ monocytes reprogrammed by exposure to either M-CSF, GM-CSF or GMCSF + IL-4 have been extensively studied as *in vitro* models for macrophages, “M2-like” macrophages or DCs (Akagawa et al., 2006; Randolph et al., 2008; Sallusto and Lanzavecchia, 1994; Xue et al., 2014). For example, to model DCs, moDCs (Sallusto and Lanzavecchia, 1994) were used. In the past, using these *in vitro* generated cells as models for macrophages and DCs was justified by morphological, phenotypical and functional similarities to *in vivo* found cellular subsets. However, recently it became clear that macrophages, monocytes and DCs present with very different transcriptomic make ups *in vivo*, reflecting their different ontogeny (Geissmann et al., 2010; Guilliams et al., 2014; Merad et al., 2013; Schlitzer et al., 2015a). Therefore, it is imperative to globally reassess the relationships between M-CSF, GM-CSF and GM-CSF + IL-4 reprogrammed monocytes and their alleged *in vivo* counterparts on a transcriptomic, phenotypic and functional level.

By using public data of ovarian cancer ascites associated infM and infDC (Segura et al., 2013), we revealed that M-CSF reprogrammed monocytes aligned closely to infM on the transcriptional level. This was phenotypically supported by common low expressions of CD1a and FcsR1. In addition, Mo-GMCSF^IL4^ aligned closely to infDCs, with common phenotypic high expressions of CD1a, CD206 alongside with FcsR1. Taken together, the currently available *in vitro* models best resemble inflammatory rather than homeostatic DC and macrophage populations, therefore, serving best as reductionist models to study the role of these cells in inflammation. For the future, we encourage to identify new culture conditions for monocyte-derived cells that resemble homeostatic DC or macrophage phenotypes guided by transcriptomic evaluation, as shown for *in vitro* cultures of microglia (Butovsky et al., 2013), and the identification of dedicated progenitors of DCs in the human blood and bone marrow (Breton et al., 2015; Lee et al., 2015).

Genome-wide assessments, as exemplified here on transcriptome level, allowed us to define the cellular relationships between these different model systems on a much more precise level and revealed an unexpectedly close association of monocytes differentiated by MCSF and GMCSF, while IL4 was the major driver for moDC identity. In addition, we found that monocytes integrate the GMCSF and IL4 signals over time, which necessitates a reassessment of our dichotomous definition of monocytes differentiating towards a macrophage or DC-like phenotype. Clearly, the varying time of onset and the variance in overall exposure to IL4 resulted in gradually changing transcriptional and functional identities of monocyte-derived cells. The unexpectedly high heterogeneity unravelled by applying high-dimensional phenotyping by mass cytometry added a further perspective of individual reactivity to the moDC response repertoire. These observations challenge our view of a strict differentiation switch between monocytes with macrophage or DC functionality induced by a single cytokine, but rather support a more dynamic differentiation model, in which cell identity is a function of nature and the duration of this input. This further corroborates that these monocyte-derived *in vitro* cultures model the more plastic cell types originating from monocytes during inflammation *in vivo* - such as infM and infDCs – than steady state macrophages or DCs.

Using our global transcriptional approach, we identified the co-repressor NCOR2 as one of the key transcriptional hubs linked to IL4-dependent differentiation of monocytes, which has not been described so far. Co-repressors such as NCOR2 play an important role throughout development and for homeostatic processes in muscle, adipose, and liver tissues (Mottis et al., 2013). NCOR2 has been shown to be important in cell fate determination, cell differentiation and lineage progression in the nervous system (Jepsen et al., 2007), and elevated expression of NCOR2 was mainly observed in tissues with high OXPHOS activity, similar to our observations of elevated expression of NCOR2 in Mo-GMCSF^IL4^ (Reilly et al., 2010). Additionally, via signalling by nuclear receptors such as peroxisome proliferator-activated receptor-γ (PPAR-γ) or liver X receptor (LXR), NCOR2 has been linked to the repression of NF-kB target genes in response to LPS stimulation of macrophages (Hoberg et al., 2004; Pascual et al., 2005). Nevertheless, a link from IL4-mediated signalling to NCOR2 remains elusive. Although it has been speculated that IL4 activation of human monocytes leads to endogenous PPAR-γ ligand production (Czimmerer et al., 2012), further work is necessary to establish a link to NCOR2-mediated gene repression. Since regulation of co-repressors such as NCOR2 are tissue-, cell type- and context-dependent (Mottis et al., 2013), future studies will pinpoint, whether transcriptional control, alternative splicing, post-translational modifications, cellular localization of NCOR2, or intermediate signalling cascades or a combination thereof are involved in the molecular function of NCOR2 in IL-4IL4- driven monocyte differentiation.

In addition, we revealed unexpected heterogeneity within the monocyte cultures and finally introduce a novel set of surface markers with superior resolution on differentially stimulated monocyte cell subsets when compared to individual markers. In particular, within Mo-GMCSF^IL4^ we identified subsets of cells that either expressed HLA-DR and CD86 or CD1a and FcεR1, the former representing a subpopulation with already elevated antigen presenting capacity. While induction of FcεR1 on CD1a^+^ DCs derived from CD34^+^ stem cells has been reported (Allam et al., 2004; Magerstaedt et al., 1997), this has not been studied during monocyte to Mo-GMCSF^IL4^ differentiation. Therefore, the heterogeneity of this event was not yet appreciated. Considering findings that CD1a^+^ and CD1a^−^ Mo-GMCSF^IL4^ differ in their capacity to direct Th cell differentiation (Cernadas et al., 2009; Chang et al., 2000; Gogolak et al., 2007) a better monitoring of these cultures in a clinical setting might be beneficial for optimizing efficiency of cellular vaccines in the future. Consequently, high-dimensional characterization should be used to further optimize culture conditions, generate more homogenous cell populations, and therefore open novel avenues for optimizing cellular products derived from human monocytes for vaccination strategies.

## Acknowledgement

We thank all members of the Schultze and Schlitzer Labs for critically reading the manuscript. In addition, we thank Johannes Oldenburg for providing us with buffy coats from healthy individuals. This work was supported by the German Research Foundation (SFB 832, SFB 704, INST 217/577-1 and SFB 738 (CF)), core grants of the Singapore Immunology Network (FG, EWN) and the Singapore Immunology Network immune-monitoring platform funding (EWN). JLS, AS, WK, EL, SB, and MB are members of the Excellence Cluster ImmunoSensation. The research leading to these results has received funding from the People Program (Marie Curie Actions) of the European Union’s Seventh Framework Program FP7/2077-2013 under REA grant agreement no. 317445. AS is supported by an Emmy Noether fellowship of the German research foundation (SCHL 2116/1-1) and an A*STAR BMRC Young Investigator Award. NMcG is supported by a Royal Society and Wellcome Trust grant (204464/Z/16/Z). We thank M. Ballmaier for cell sorting at MHH.

## Material and Methods

### Ethics statement

Buffy coats were obtained from healthy donors in cooperation with the University hospital Bonn (local ethics vote 203/09) after written consent was given according to the Declaration of Helsinki.

### Isolation of human mononuclear cells from buffy coats

Peripheral blood mononuclear cells (PBMC) were isolated by Pancoll (PAN-Biotech) density centrifugation from buffy coats. CD14-, CD56-, CD4- and CD19-specific MACS beads (Miltenyi Biotec) were used for the enrichment of CD14^+^ monocytes, CD56^+^ NK cells, CD4^+^ T cells and CD19^+^ B cells, respectively. Lung-derived monocytes were isolated from human perfusates of lung transplant recipients with informed consent and immediately sorted for CD45^+^ CD14^+^HLA-DR^high^ cells using the FACS Fusion cell sorter (BD, USA).

### Generation of reprogrammed CD14^+^ monocytes

CD14^+^ monocytes were cultured in RPMI1640 medium supplemented with 10% FCS, 1% Penicillin-Streptomycin, 1% Sodium pyruvate and 1% Glutamax (all from Gibco) for three days. CD14^+^ monocytes were differentiated into Mo-MCSF or Mo-GMCSF in the presence of 100 IU/ml rhMCSF or 800 IU/ml rhGMCSF, respectively. Mo-GMCSF^IL4^ were generated by the addition of 800 IU/ml rhGMCSF and 500 IU/ml rhIL4 and were incubated for up to 6 days. All cytokines were purchased from Immunotools.

### Flow cytometry

Cells were washed with ice cold PBS. After FcR blockage (Miltenyi, Germany), cells were stained with the respective antibodies in PBS supplemented with 0.5% FCS, 2.5 mM EDTA for 20min at 4°C. Following antibodies were purchased from Biolegend (USA): CD19 (HIB19), CD56 (HCD56), CD11b (ICRF44), CD23 (EBVCS-5), HLA-DR (L243), CD14 (M5E2); R&D: VSIG4 (polyclonal), CD1E (704407), MARCO (polyclonal); Becton Dickinson (BD, USA): CD209 (DCN46), CD226 (DX11); Data acquisition was performed with a LSR II (BD). Analyses were performed with FlowJo software (Tree Star).

### Mass cytometry

Following culture, cells were washed with PBS (GIBCO, Life Technologies, Carlsbad, CA) and stained with cisplatin (Sigma-Aldrich, St Louis, MO) on ice to exclude dead cells. Afterwards, cells were washed with PBS containing 4% FBS and 0.05% Sodium azide and fixed with 2% paraformaldehyde (PFA; Electron Microscopy Sciences, Hatfield, PA) overnight. Cells were permeabilized (1X perm buffer (Biolegend, San Diego, CA)) and stained with metal-conjugated antibodies (**Table S7**) intracellularly. Then cells were washed, metal barcoded and resuspended in water. EQ Four Element Calibration Beads (Fluidigm Corporation, South San Francisco, CA) were added at a concentration of 1% prior to acquisition. Cells were acquired and analyzed using a CyTOF1 Mass cytometer. The data was normalized (Finck et al., 2013) and events with parameters having zero values were replaced with a randomized value between 0 and −1. Cells for each barcode were deconvoluted (using manual gating to select cells stained with two and only two barcoding channels.

### Mass cytometry data analysis

Normalized mass cytometry data was exported in .fcs format and pre-processed in FlowJo Version 9.9.4 (Tree Star Inc). Pre-processing included removal of cell debris (Iridium-191/193 DNA intercalating fraction) and dead cells (cisplatin^+^). Myeloid CD45^+^lin(CD3/CD7/CD15/CD19/CD20/CD56)^−^ cells were exported and used for analysis. Downstream analysis was performed using the Cytofkit R package. For comparison of the different reprogramming conditions data from 1000 myeloid cells was randomly sampled per donor and condition (3 donors; 12,000 cells in total) and autoLgcl-transformed including expression values for 36 surface markers (CD45, CD14, CD5, CD62L, CD48, CD68, CD66ace, CLA, HLA-DR, CD115, CD64, CD1c, FcεR1, SEPP1, CD123, CD163, CXCR3, CD226, CD169, SIRP1a, Dectin1a, CD1a, CD141, MARCO, CD86, CX3CR1, CD206, VSIG4, CD88, CD34, MerTK, CD39, CD26, CD11c, CD11b, CD16). Detailed analysis on the Mo-GMCSF^IL4^ condition was based on 3500 cells from one individual. To define clusters of cell subpopulations, PhenoGraph method was used. Points representing individual cells in the *t*-SNE plots were color-coded to illustrate levels of protein expression or affiliations to clusters, treatment conditions or donors, respectively. Alternatively, the gplots R package was used to generate heatmaps of marker expression of individual cells or mean values over identified cell clusters. Dendrograms representing hierarchical clustering results based on the Euclidean distance measure also were included.

### Uptake of fluorescent microbeads or yeast

Cells were incubated either with fluorescent monodispersed polystyrene microspheres (1 μm diameter, Fluoresbrite YG Beads, Polysciences) or yeast (GFP-expressing *Pichia Pastoris*) in a cell-to-bead ratio of 1/10 for 4h or 60 min at 37°C, respectively (Kreer et al., 2017). Afterwards, cells were harvested, washed and bead/yeast uptake was analyzed by flow cytometry using an LSR II (BD). Negative control samples were kept at 4°C. Data analysis was performed using FlowJo software (Tree Star).

### Western blot

For hNCOR2 protein detection, the cytosolic and nuclear whole protein fractions were separated by SDS-PAGE and transferred onto a nitrocellulose membrane (Amersham) by wet blotting. Probing was performed using hNCOR2 antibody (Abcam). Signal expression values were calculated in semi-quantitative relation to HSP70 (BioTrend) and β-actin expression values (Millipore). For MMP12 protein detection, the cytosolic whole protein fractions were separated by SDS-PAGE and transferred onto a nitrocellulose membrane (Amersham) by semi-dry blotting. Probing was performed by using MMP-12 antibody (R&D). Signal expression values were calculated in semi-quantitative relation to β-actin expression values (Millipore). All signal detection and analysis was performed via the LI-COR Odyssey system. Dignam extraction efficiency was validated for enrichment of cytosolic proteins in the cytosolic fractions compared to the nuclear proteins in nuclear fractions via HDAC1 (Cell signaling) versus GAPDH (Cell signaling) detection.

### Migration assay

Migration was analyzed in μ-Slide 8 well chambered coverslips (Ibidi) coated with 50μg/ml human fibronectin (Alfa Aesar). 0.7×10^5^ cells in 300μl VLE-RPMI (Biochrom) were transferred to each well. Live cell imaging of adherent cells was performed at 37°C and 5% CO_2_ using a fully automated inverted Olympus Fluoview 1000 confocal microscope equipped with motorized xyz stage (Marzhauser). Cell motility was monitored over a period of 3h by capture of differential interference contrast images every 5min with a 0.40 UPLAPO 10× Objective (Olympus). Migration parameters were calculated using the Manual Tracking and Chemotaxis Tool plugins in ImageJ.

### Nanostraw-facilitated NCOR2 siRNA knockdown

24h prior to the experiment, the nanostraw cargo chamber was washed three times with 10μL of 0.5% PEG 3500 (P3640, sigma) in PBS and equilibrated; chambers were equilibrated with 100μL RPMI 1640 media. CD14^+^ human monocytes were resuspended in RPMI 1640 with supplements (10% FCS, 1% Pen/Step, 1% GlutaMax and 1% NaPyruvat) and activated. siRNA solutions were prepared in 1× siRNA Buffer (Dharmacon). hNCOR2 siRNA (Dharmacon) ON Target Plus was used for knockdown. Immediately prior to the experiment, the equilibration media was removed, the tubing system was flushed with the siRNA solution or PBS only, and completely filled. Subsequently, the cell suspension was filled into the chamber and incubated for 72h at 37°C and 5%CO_2_. After 72h, cells were directly lysed within the chambers by adding Trizol. qRT-PCR was performed to check transfection efficiency.

### Cell stimulation

Mo-MCSF, Mo-GMCSF, Mo-GMCSF^IL4(0-72h)^ or Mo-GMCSF^IL4(0-144h)^ cells were stimulated overnight under the following conditions: Media, 100ng/ml LPS ultrapure, 100ng/ml LPS ultrapure + 1000U/ml IFN-γ, 1μg/ml CL097, 100ng/ml flagellin. Supernatants were harvested after 19h of stimulation and stored at −80°C until further processing for cytokine analysis.

### Cytokine measurement

Cytokines were measured using LEGENDplex. Briefly, diluted cell culture supernatants were incubated for 2 hours with the provided beads and detection antibodies, followed by another 30min incubation with SA-PE. After washing, beads were resuspended in washing buffer and acquired using a LSRII flow cytometer (BD). Data were analyzed with the LEGENDplex Data Analysis Software; concentration values were exported to Excel and visualized in R.

### Oxygen consumption rate (OCR) and ECAR measurements

OCR and ECAR were determined using a XF-96 Extracellular Flux Analyzer (Seahorse Bioscience). For ECAR analysis the media was changed to bicarbonate-free RPMI supplemented 10mM glucose, 1mM pyruvate & 2 mM glutamine 1h prior to the assay and the plate was kept in a non-carbonated incubator. Measurements were performed under basal conditions and after the sequential addition of final 1μM oligomycin A, 1.5μM FCCP (fluoro-carbonyl cyanide phenylhydrazone) and 0.5μM rotenone & antimycin each. For ECAR analysis the media was changed to bicarbonate free RPMI supplemented 2 mM glutamine 1h prior to the assay and the plate was kept in a non-carbonated incubator. Measurements were performed under basal conditions and after the sequential addition of final 10 mM glucose, 1 μM oligomycin and 100 mM 2-Deoxyglucose. OXPHOS was calculated as (basal OCR – OCR after rotenone & antimycin treatment), ATP production was calculated as (basal OCR – OCR after oligomycin A treatment), maximal respiration was calculated as (OCR after FCCP treatment – OCR after rotenone & antimycin treatment), glycolysis was calculated as (basal ECAR – ECAR after 2-Deoxyglucose treatment), glycolytic capacity was calculated as (ECAR after oligomycin A treatment - ECAR after 2-Deoxyglucose treatment. All reagents were purchased from Sigma, except FCCP was purchased from Tocris. ECAR and OCR raw data was normalized to DNA content using the CyQuant Assay kit (Thermo Fisher).

### Microarray data generation

Up to 5 × 10^6^ cells were harvested and lysed in TRIzol (Invitrogen) and RNA was isolated and concentration and purity was assessed using a NanoDrop 1000 UV-Vis Spectrophotometer (Thermo Scientific). Subsequently, the TargetAmp-Nano Labeling Kit for Illumina Expression BeadChip (Epicentre) was utilized to generate biotin labeled anti-sense RNA (cRNA) according to the manufacturer’s protocol. As a quality control, 100 ng cRNA were reverse transcribed to cDNA and a PCR specific for *ACTB* amplification was performed. For expression profiling, 750 ng cRNA were hybridized to Human HT-12v3 BeadChip arrays (Illumina), stained and imaged on an Illumina iScan system.

### RNA-sequencing

Total RNA was converted into libraries of double stranded cDNA molecules as a template for high throughput sequencing following the manufacturer’s recommendations using the Illumina TruSeq RNA Sample Preparation Kit v2. Shortly, mRNA was purified from 100 ng of total RNA using poly-T oligo-attached magnetic beads. Fragmentation was carried out using divalent cations under elevated temperature in Illumina proprietary fragmentation buffer. First strand cDNA was synthesized using random oligonucleotides and SuperScript II. Second strand cDNA synthesis was subsequently performed using DNA Polymerase I and RNase H. Remaining overhangs were converted into blunt ends via exonuclease/polymerase activities and enzymes were removed. After adenylation of 3′ ends of DNA fragments, Illumina PE adapter oligonucleotides were ligated to prepare for hybridization. DNA fragments with ligated adapter molecules were selectively enriched using Illumina PCR primer PE1.0 and PE2.0 in a 15 cycle PCR reaction. Size-selection and purification of cDNA fragments with preferentially 200 bp in length was performed using SPRIBeads (Beckman-Coulter). Size-distribution of cDNA libraries was measured using the Agilent high sensitivity DNA assay on a Bioanalyzer 2100 system (Agilent). cDNA libraries were quantified using KAPA Library Quantification Kits (Kapa Biosystems). After cluster generation on a cBot, a 75 bp single-read run was performed on a HiSeq1500.

### Summary of bioinformatic analyses of microarray and RNA-Seq data

Detailed descriptions of all bioinformatic analyses can be found in the Supplemental Information. Briefly, PCAs, PCCMs, and heatmaps of the 1000 most variable probes or genes were used to investigate the relationships between samples of the different datasets. Linear SVR (Newman et al., 2015), GSEA (Subramanian et al., 2005) and correlation analyses were applied to study the transcriptional similarities between Mo-GMCSF^IL4 (0-72/144h)^ cells and infDCs, as well as between Mo-MCSF, Mo-GMCSF and infM (GSE40484, (Segura et al., 2013)). Genes being either commonly expressed in all monocyte-derived cells, but not in CD14^+^ monocytes, or in only a specific monocyte-derived subset, were determined based on differential expression analyses combined with filtering. Co-expression networks were generated either based on unions of differentially expressed genes or present TRs. GOEA was performed on genes being up- or downregulated in Mo-GMCSF in direct comparison to Mo-MCSF. Heatmaps of genes were created, which were linked to explored functions (migration, bead uptake and OCR/ECAR), and which recapitulated the functional outcomes in terms of magnitude. SOM Clustering was performed on all genes being present in the RNA-Seq time kinetics dataset. IL4-related gene signatures were established by comparing IL4 treated to IL4 untreated myeloid cells of the three public datasets GSE13762 (Széles et al., 2009), GSE35304 (Clayberger et al., 2012) and GSE32164 (Pello et al., 2012). These signature were compared to genes being variable between siRNA anti-NCOR2 and WT RNA-Seq data samples. Microarray and RNA-Seq data were uploaded to the Gene Expression Omnibus database (www.ncbi.nlm.nih.gov/gds) and can be found under the accession number GSE96719. These signatures were compared to genes being variable between siRNA anti-NCOR2 and WT RNA-Seq data samples. Microarray and RNA-Seq data were uploaded to the Gene Expression Omnibus database (www.ncbi.nlm.nih.gov/gds) and can be found under the accession number GSE96719.

---

**Supplementary figure 1.**
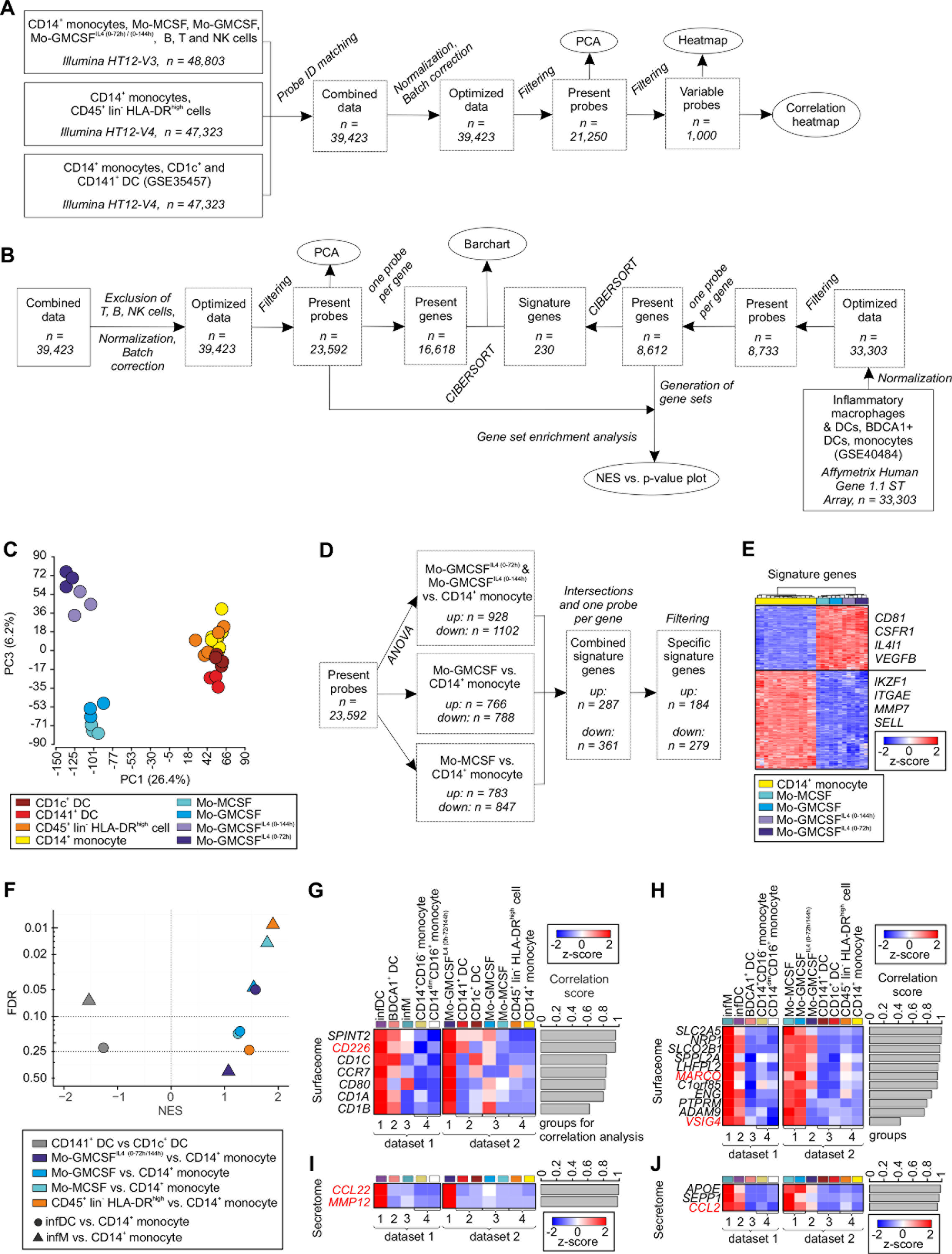

**Supplementary figure 2.**
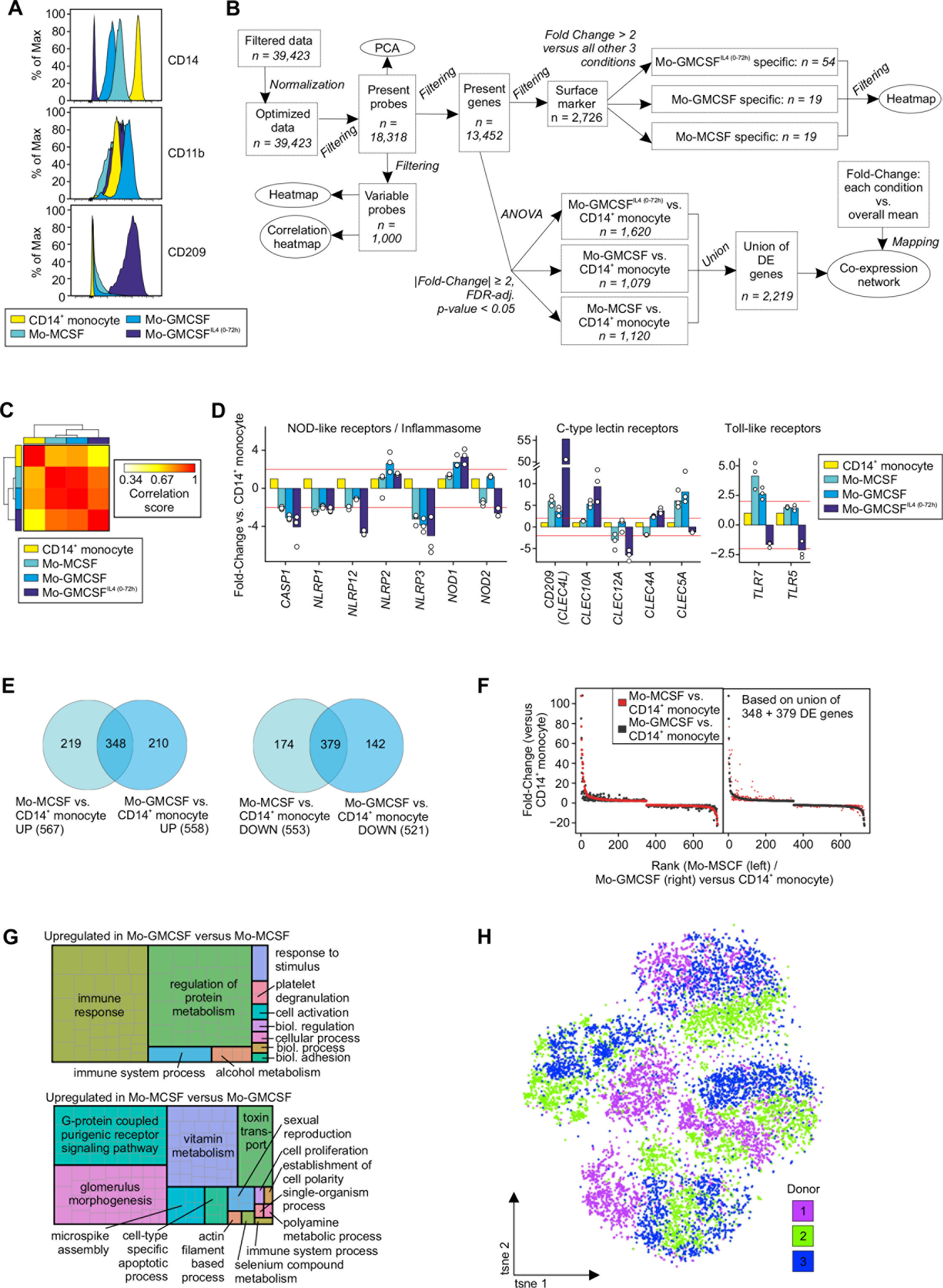

**Supplementary figure 3.**
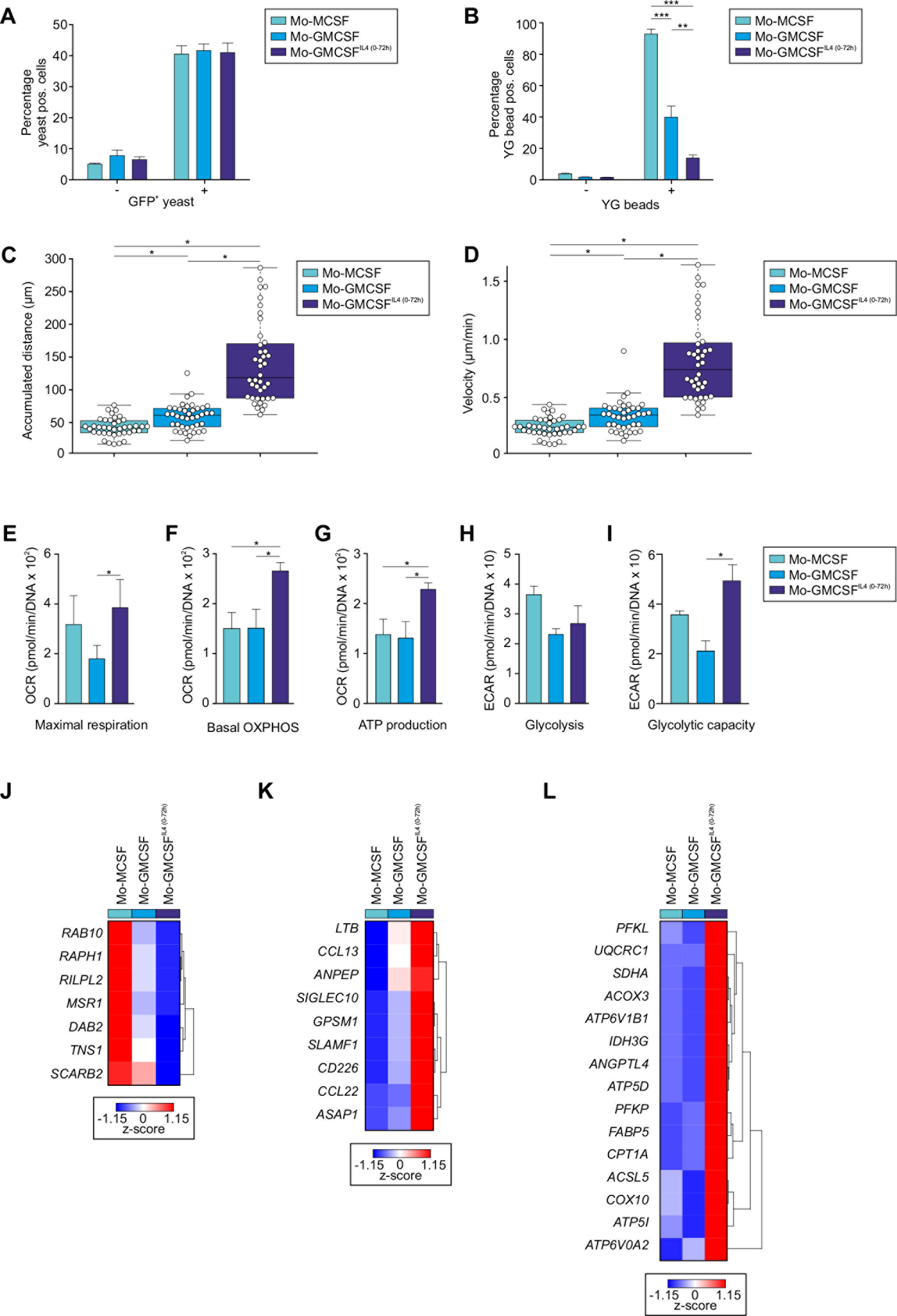

**Supplementary figure 4.**
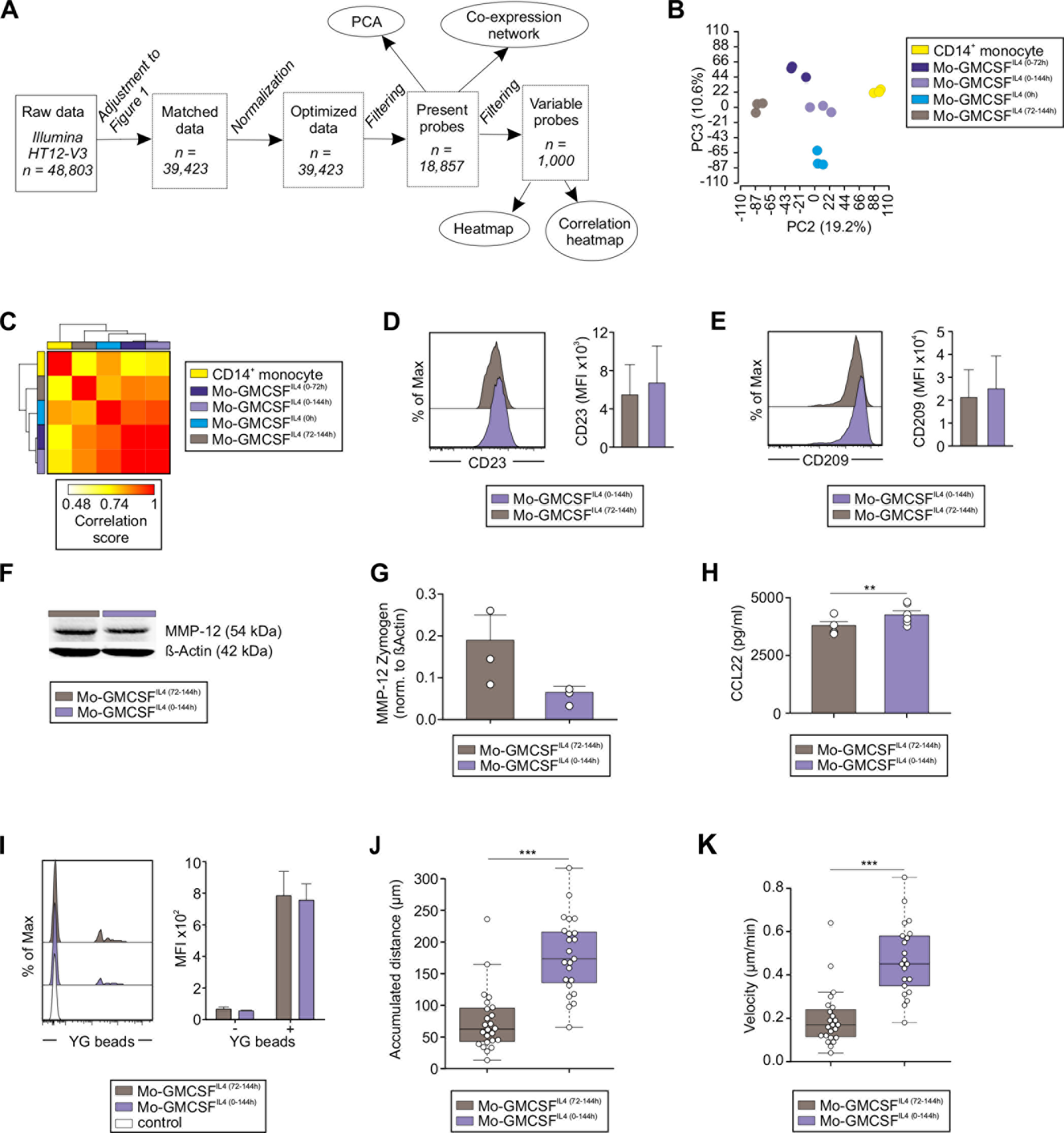

**Supplementary figure 5.**
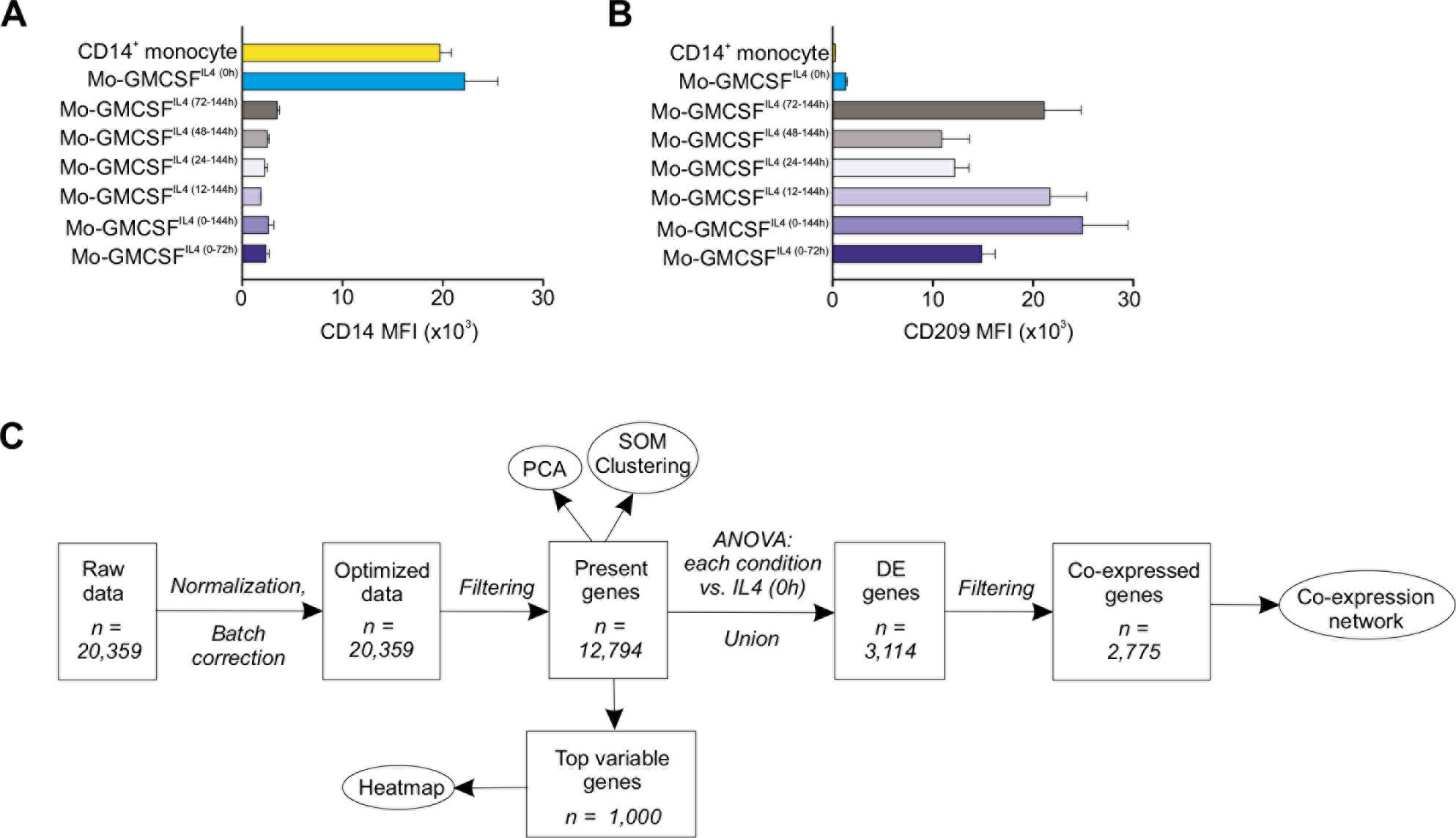

**Supplementary figure 6.**
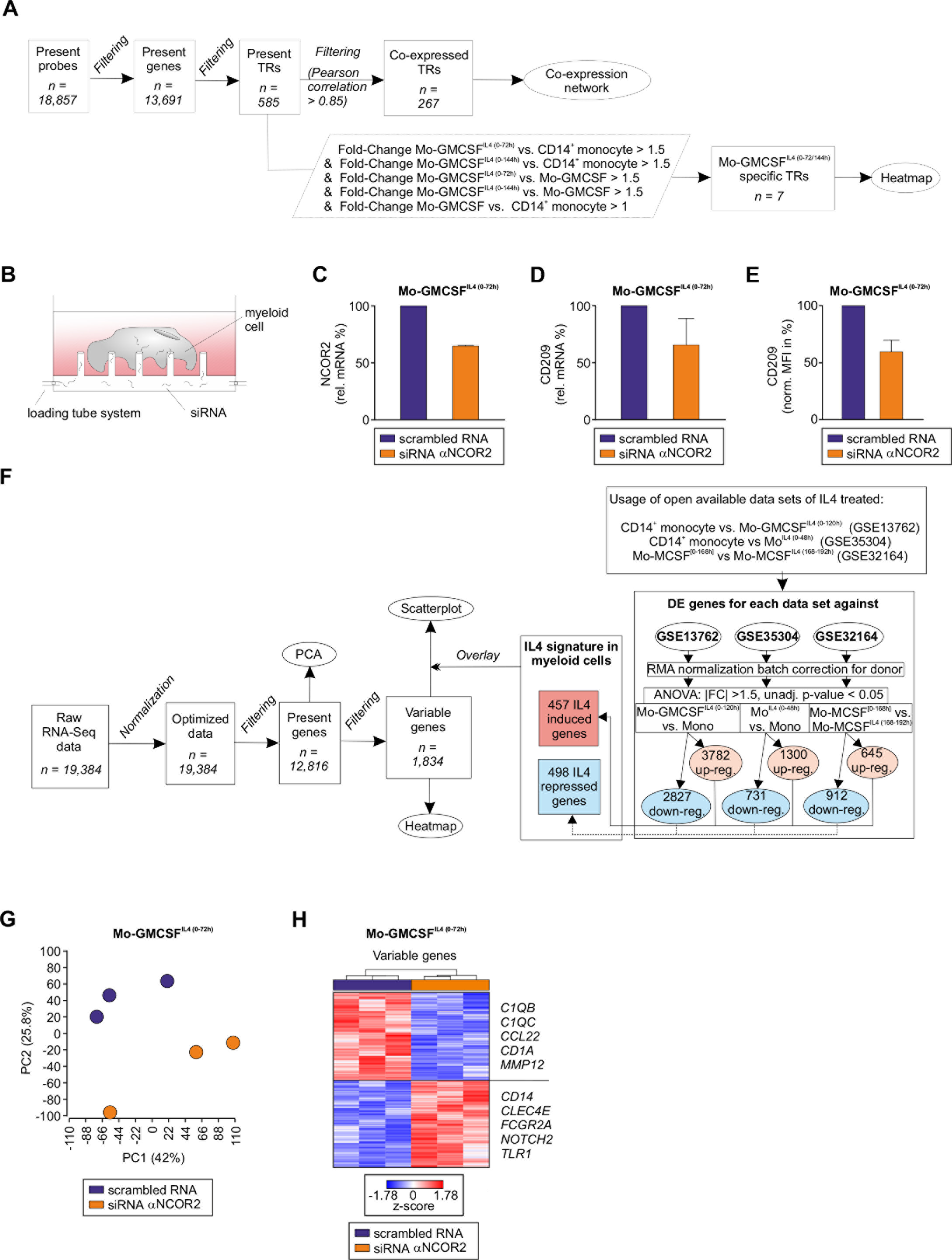

